# Alpha-1 Antitrypsin Limits Neutrophil Extracellular Trap Disruption of Airway Epithelial Barrier Function

**DOI:** 10.1101/2022.03.18.484920

**Authors:** KM Hudock, MS Collins, M Imbrogno, EL Kramer, JJ Brewington, A Ziady, N Zhang, J Snowball, Y Xu, BC Carey, Y Horio, SM O’Grady, EJ Kopras, J Meeker, H Morgan, AJ Ostmann, E Skala, ME Siefert, CL Na, K Gollomp, N Mangalmurti, BC Trapnell, JP Clancy

## Abstract

Neutrophil extracellular traps contribute to lung injury in cystic fibrosis and asthma, but the mechanisms are poorly understood. We sought to understand the impact of human NETs on barrier function in primary human bronchial epithelial and a human airway epithelial cell line. We demonstrate that NETs disrupt airway epithelial barrier function by decreasing transepithelial electrical resistance and increasing paracellular flux, partially by NET-induced airway cell apoptosis. NETs selectively impact the expression of tight junction genes claudins 4, 8 and 11. Bronchial epithelia exposed to NETs demonstrate visible gaps in E-cadherin staining, a decrease in full-length E-cadherin protein and the appearance of cleaved E-cadherin peptides. Pretreatment of NETs with alpha-1 antitrypsin (A1AT) inhibits NET serine protease activity, limits E-cadherin cleavage, decreases bronchial cell apoptosis and preserves epithelial integrity. In conclusion, NETs disrupt human airway epithelial barrier function through bronchial cell death and degradation of E-cadherin, which are limited by exogenous A1AT.

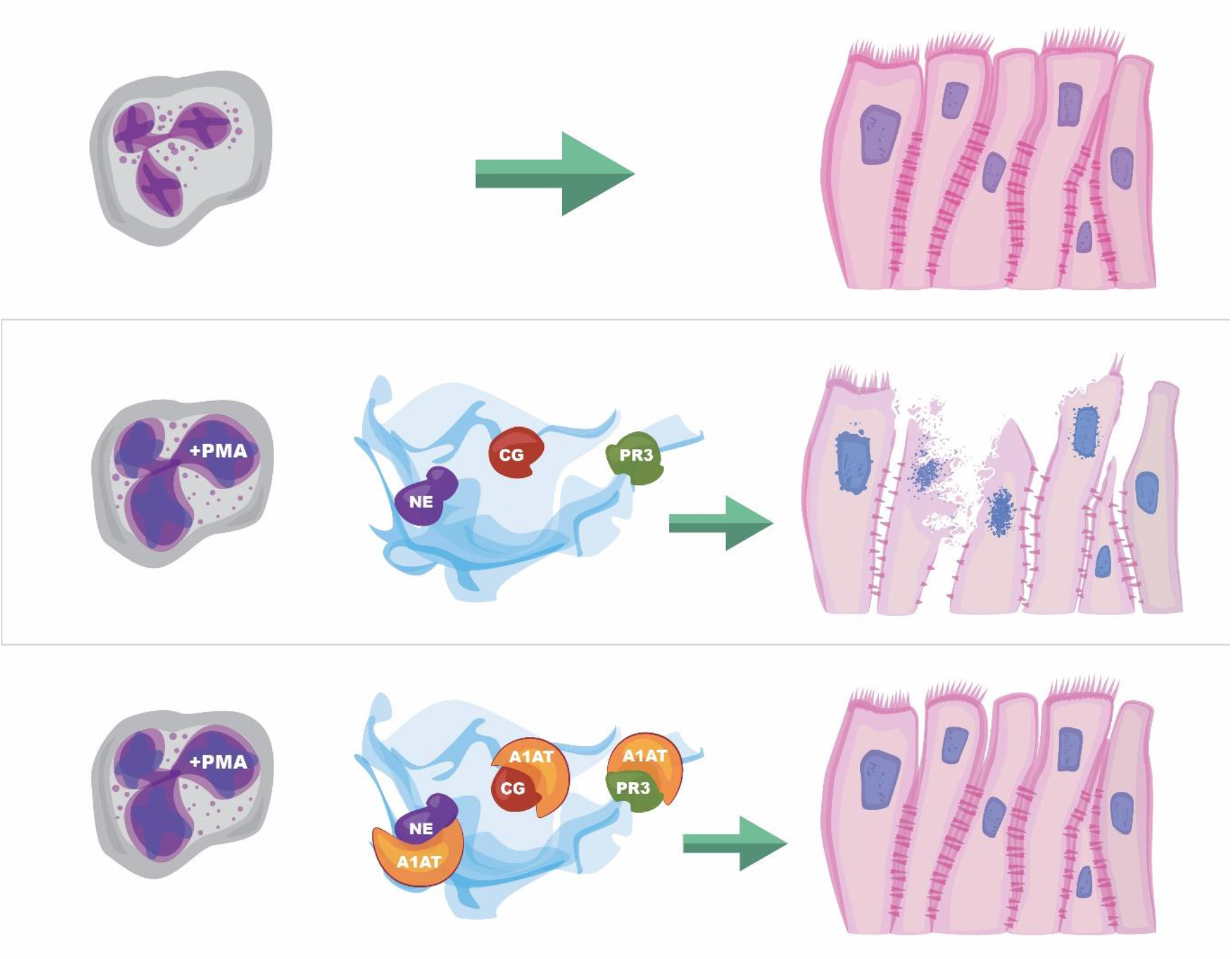

## Introduction

Neutrophils play a critical role in innate host defense. Neutrophil extracellular traps (NETs) are web-like structures released by neutrophils in response to numerous pathogens, inflammatory cytokines and other compounds (Young et al., 2011, Veras et al., 2020, Keshari et al., 2012, Kenny et al., 2017). NETs trap bacteria in the bloodstream and tissues to limit dissemination to other organs (McDonald et al., 2012, Thanabalasuriar et al., 2019, Gollomp et al., 2020). NETs also play a role in thrombus formation via multiple mechanisms (Gollomp et al., 2018, Fuchs, T. A. et al., 2010, Shi et al., 2021). NETs are comprised of a DNA backbone studded with histones, proteases, cytokines and other immunomodulatory molecules (Brinkmann et al., 2004). The most commonly reported NET proteases are neutrophil elastase (NE) and cathepsin G (CG) (Chapman et al., 2019). NET proteases are enzymatically active even when exposed to recombinant human DNase (Hudock et al., 2020). NET proteases perform functions such as processing of cytokines into active forms and degrading bacterial virulence factors (Clancy et al., 2018, Brinkmann et al., 2004). NETs have been implicated in perpetuating airway inflammation in conditions including asthma and CF and we have previously reported that NETs can increase IL-8 via selective activation of the IL-1 pathway (Lachowicz-Scroggins et al., 2019, Manzenreiter et al., 2012, Hudock et al., 2020). The impact of NETs on other critical epithelial roles such as barrier function, however, remains largely unstudied.

Bronchial epithelia serve an important role in barrier function to limit entry of pathogens and injurious agents into the submucosa and bloodstream. Human bronchial airways are predominately lined with ciliated pseudostratified columnar epithelial cells and an overlying thin layer of mucus (Whitsett, 2018). Epithelial cells form an effective barrier through a complex junctional protein network that maintains cell-cell contact and limits paracellular passage (Montefort et al., 1993). Junctional protein structures include tight junctions, adherens junctions, and desmosomes. E-cadherin is a crucial adherens junctional protein that is widely expressed in human epithelia and is a key component of the intact human airway mucosa (Nawijn et al., 2011). Respiratory infections, e.g., adenovirus and *Pseudomonas aeruginosa*, are associated with loss of E-cadherin (Man et al., 2000, Reboud et al., 2017). Decreased E-cadherin compromising epithelial barrier function has been reported in airway diseases including asthma (Nawijn et al., 2011). The neutrophil protease MMP-9 enhanced conductance of human airway epithelia by altering localization of the tight junction protein claudin-1. (Vermeer et al., 2009) In addition to disrupted barrier function, alterations in epithelial junctional molecules have profound consequences for cell-cell signaling, cellular differentiation and spread of malignant metastases (Nawijn et al., 2011, Brune et al., 2015).

Alpha-1 antitrypsin (A1AT) is a naturally occurring serine protease inhibitor that binds to and mitigates the activity of serine proteases including NE, CG and proteinase 3 (PR3). A1AT is predominately generated by hepatocytes, but human bronchial epithelia and neutrophils produce A1AT as well (Cichy, Potempa & Travis, 1997, du Bois et al., 1991, Clemmensen et al., 2011). A1AT deficiency is due to a protease-antiprotease imbalance that results in unhindered protease activity, increased inflammation and destruction of lung tissue (McCarthy, Reeves & McElvaney, 2016). Although A1AT inhibits neutrophil proteases, it is unclear if it effectively binds NET proteases and limits lung epithelial injury.

In the current report we hypothesized that NETs were sufficient to alter airway epithelial barrier function. To test this hypothesis, we exposed primary human bronchial epithelia (HBE) grown at air liquid interface (ALI) and a polarized human bronchial epithelial cell line (wtCFBE41o-) to isolated human NETs *in vitro*. Our work demonstrated that NETs disrupted human airway epithelia barrier function assessed by electrical resistance, transepithelial macromolecule flux, confocal microscopy, transmission electron microscopy, and junctional RNA and protein expression. Several NET-driven mechanisms contributed to loss of barrier function, including induction of apoptosis and degradation of E-cadherin. A1AT partially limited NET-driven cleavage of E-cadherin, bronchial cell apoptosis and maintained monolayer resistance. Together, our results demonstrate that NETs produce airway cell injury and death leading to disruption of bronchial epithelial integrity. This study links NET exposure to loss of epithelial cell integrity, which may drive morbidity and mortality in many lung diseases. Importantly, we also show a potential role for A1AT, an FDA approved drug, in ameliorating these detrimental NET effects.

## Results

### Human NETs disrupt epithelial monolayer integrity

We sought to determine if isolated human NETs disrupt barrier function of HBE *in vitro*. Neutrophils were harvested from the peripheral blood of healthy human donors using negative bead selection and exposed to phorbol 12-myristate 13-acetate (PMA) to stimulate NETosis (Najmeh et al., 2015, Hudock et al., 2020). As previously described, the resulting viscous layer of NETs was collected and centrifuged to remove cellular debris (Najmeh et al., 2015). Primary HBE grown at ALI or wtCFBE41o-cell line expressing wild type CFTR were exposed to isolated NETs in the apical compartment of cell culture inserts. Transepithelial electrical resistance (TEER) was measured to assess the barrier function of the epithelial monolayer at baseline and after exposure to NETs. Both wtCFBE41o- and HBE maintain tight cell-cell junctions across the monolayer reflected by high baseline resistance measurements (686Ω/cm^2^ +/-61.12, 1238Ω/cm^2^ +/-31.62, respectively) (Haws et al., 1992, Brewington et al., 2018).

wtCFBE41o-cells exposed to NETs for 18 hours had a ∼50% decrease in electrical resistance when compared to media control, which was unchanged (Figure 1A). Primary HBE exposed to NETs had a similar average reduction (Figure 1B). Pretreatment of NETs with dornase alpha (DA), a DNase therapy prescribed for CF, did not limit NET-driven reductions in resistance (Fuchs, H. J. et al., 1994). DA alone yielded similar results to controls in both HBE and wtCFBE41o-. Resistance measured in primary HBE after 2 and 6 hour NET exposure were unchanged compared to PBS, but significantly decreased at 18 hours (p<0.0001) (Supplemental Figure 1A). To test if individual components of NETs recapitulated NET reductions in electrical resistance, HBE were exposed to NETs, heat inactivated NETs (to disrupt enzymatic activity) or individual NET proteases compared to PBS. All of these significantly decreased TEER compared to PBS. None of these decreased TEER at 18 hours to the extent seen with NETs (Supplemental Figure1B). Exposing HBE for 18 hours to a mix of NET components composed of three serine proteases, gDNA and citrullinated histones in similar concentrations as found in intact NETs did not recapitulate the decrease in TEER seen with NETs (Supplemental Figure 1C).

**Figure 1.**
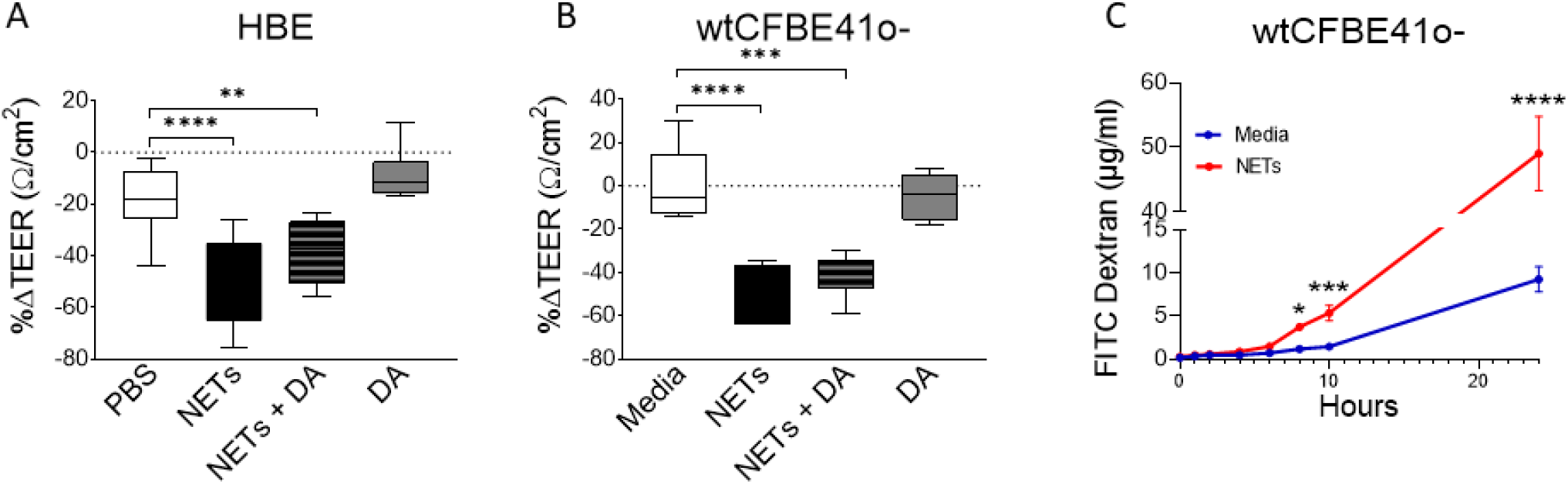
Neutrophil extracellular traps (NETs) disrupt epithelial barrier function. **(A)** wtCFBE41o-grown to confluence and **(B)** primary normal human bronchial epithelia (HBE) grown at air-liquid interface (ALI) were exposed to media or PBS control (respectively), 5μg/ml NETs, NETs + 0.5μg/ml dornase alpha (DA), or DA in the apical compartment for 18 hours in triplicate wells. Transepithelial electrical resistance (TEER) was measured pre and post treatment in duplicate for each well and expressed as a percent change. Data analyzed by one-way ANOVA followed by Bonferroni’s multiple comparisons test (experiments were performed in triplicate, HBE donors=3, NET donors=4). **(C)** Apical to basolateral flux of FITC labeled dextran across wtCFBE41o-exposed to media control or NETs from 0-24 hours. Data analyzed by Least Squares Means, F-test=264.58, ****p<0.0001 from 8-24 hours (experiments were performed in duplicate, NET donors=2). *p<0.05, **p<0.01, ***p<0.001, ****p<0.0001

To determine if the NET-driven reduction in electrical resistance was associated with altered permeability of the epithelia to large molecules, we monitored the impact of NET exposure on the apical-to-basolateral flux of FITC-labeled dextran across wtCFBE41o-monolayers (Zaidman et al., 2017). FITC-dextran concentrations increased significantly in the basolateral media beginning at 8 hours by 68.5% (media control = 0.9-1.4μg/ml vs NETs = 3.4-3.9μg/ml, p=0.0044) through 24 hours by 81.2% (media control = 7.2-11.6μg/ml vs NETs = 41.4-53.0μg/ml, p<0.0001) in wtCFBE41o-cells exposed to NETs (Figure 1C). There was no significant change in FITC dextran flux from 0-6 hours of NET exposure, which is consistent with the lack of change in HBE TEER from 0-6 hours (Supplemental Figure 1A). Together, our findings suggest that NET disruption of epithelial monolayer barrier function occurs over time, enhancing paracellular passage of macromolecules.

### NETs cause human airway cell death via apoptosis

To determine if changes in epithelial monolayer barrier function were due to NET-induced airway cell injury, we measured lactate dehydrogenase (LDH), an indicator of cytotoxicity (McGraw et al., 2020). LDH increased in the apical supernatant of HBE in response to NET exposure after 18 hours (Figure 2A). LDH release from HBE exposed to NETs or NETs pretreated with DA were similar. DA alone did not cause cytotoxicity to HBE cells.

**Figure 2.**
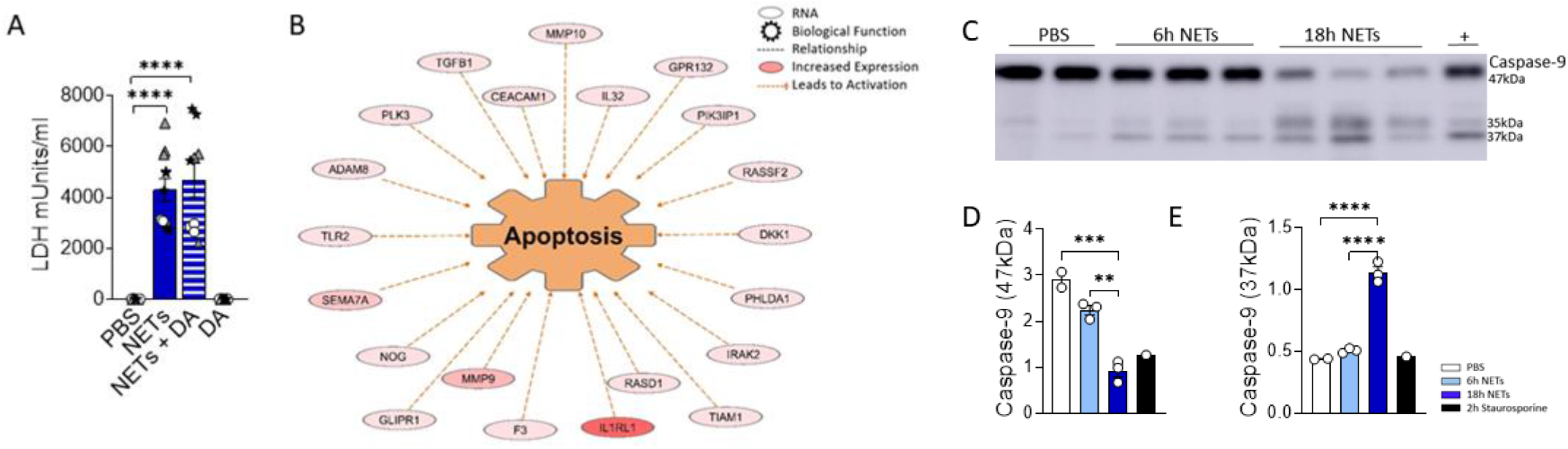
NETs activate apoptosis in normal human bronchial epithelia. HBE were grown at ALI and exposed to PBS control, 5μg/ml NETs, NETs+0.5μg/ml dornase alpha (DA), or DA in the apical compartment 18h in triplicate wells. **(A)** LDH concentration was measured in apical supernatant. Data analyzed by one-way ANOVA followed by Bonferroni’s multiple comparisons test (experiments=2, HBE donors=2, NET donors=2). **(B)** Differential gene expression assessed by RNA sequencing in HBE after exposure to 5μg/ml NETs compared to PBS. Network representation of impacted genes predicted to activate apoptosis. Color intensity indicates increased expression. Data analyzed using Ingenuity Pathway Analysis (IPA) (p=1.07e-13, experiments=3, HBE donors=3, NET donors=3). **(C)** Representative western blot of HBE lysates and analyses demonstrate **(D)** decreased full-length caspase-9, **(E)** increased cleaved caspase-9 at 37kDa, indicating apoptosis occurs in HBE exposed to NETs at 6h and 18h compared to PBS. Positive control (+) 16HBE cells treated with staurosporine (5μM, 2h). Data analyzed by one-way ANOVA followed by Bonferroni’s multiple comparisons test (experiments=2, HBE donors=2, NET donors=3). **p<0.01, ***p<0.001, ****p<0.0001

To establish which cell death pathway is triggered by NETs in human airway epithelia, we performed RNA seq on HBE from three donors exposed to NETs from 3 different individuals. A network of fifty-two RNAs involved in apoptosis had significantly increased expression and thirty RNAs had significantly decreased expression (Figure 2B, Supplemental Table 1). CAV1, which was downregulated 5-fold, encodes cavelolin, a scaffolding protein found in plasma membranes which inhibits apoptosis (Marudamuthu et al., 2019). PHLDA1 was increased 6-fold and is known to promote apoptosis via Fas expression (Park et al., 1996). RNAs for the interleukin-1 receptor and other pro-inflammatory cytokines were highly upregulated and may contribute to cytokine-driven apoptosis (Zhang, Gharaee-Kermani & Phan, 1997, Abbate et al., 2011). We also examined necroptosis, a pro-inflammatory cell death pathway that could contribute to the NET-induced inflammation we previously reported in bronchial epithelia (Hudock et al., 2020, Zhu et al., 2018). NETs had no impact on the expression of genes controlling necroptosis in epithelia after 18 hours of exposure.

To further assess the role of apoptosis in HBE exposed to NETs, cell lysates were analyzed by western blot for activation of caspase-9, an initiator of the intrinsic pathway of apoptosis. Mitochondrial release of cytochrome c, and subsequent apoptotic protease-activating factor-1 (APAF1) activation, cleaves the 47kDa procaspase-9 into active dimers detectable at 35kDa and 37kDa (Zou et al., 2003). After 6 hours of NET exposure in HBE, there was a trend towards decreasing procaspase-9 and increased cleaved caspase-9 peptides. After 18 hours procaspase-9 was significantly decreased (p=0.0005) and the cleaved caspase 9 peptides significantly increased (p=0.0139) compared to PBS (Figure 2C-D). There was no change in the necroptosis pathway proteins RIP-1, MLKL, or corresponding phosphorylated isoforms in HBE exposed to NETs (Supplemental Figure 2A-B) (Faust, Mangalmurti, 2020). NETs did not cause cleavage of pro-caspase-1 in HBE to activate pyroptosis at the measured time points (Supplemental Figure 2C). (Lee et al., 2018; Tsai et al., 2017) Together, these data support the observation that NETs activate apoptosis in primary airway epithelia.

### NETs damage epithelial cell junctions

We next investigated if NETs disrupt the airway epithelial barrier by impacting cell junctions. RNA sequencing of HBE exposed to NETs identified significant changes in expression of RNAs regulating cell junction organization (Figure 3A-3B and Supplemental Table 2). LAMC2, upregulated 8.4-fold, encodes an epithelial protein, which connects cells to the basement membrane (Salo et al., 1999). SNAI2, downregulated 4.7-fold, is thought to repress E-cadherin transcription (Reinhold et al., 2010). Along with upregulation of metalloproteinase genes, involved in extracellular matrix degradation, these transcriptional changes signal the breakdown of cell monolayer architecture (Matrisian, 1992).

**Figure 3.**
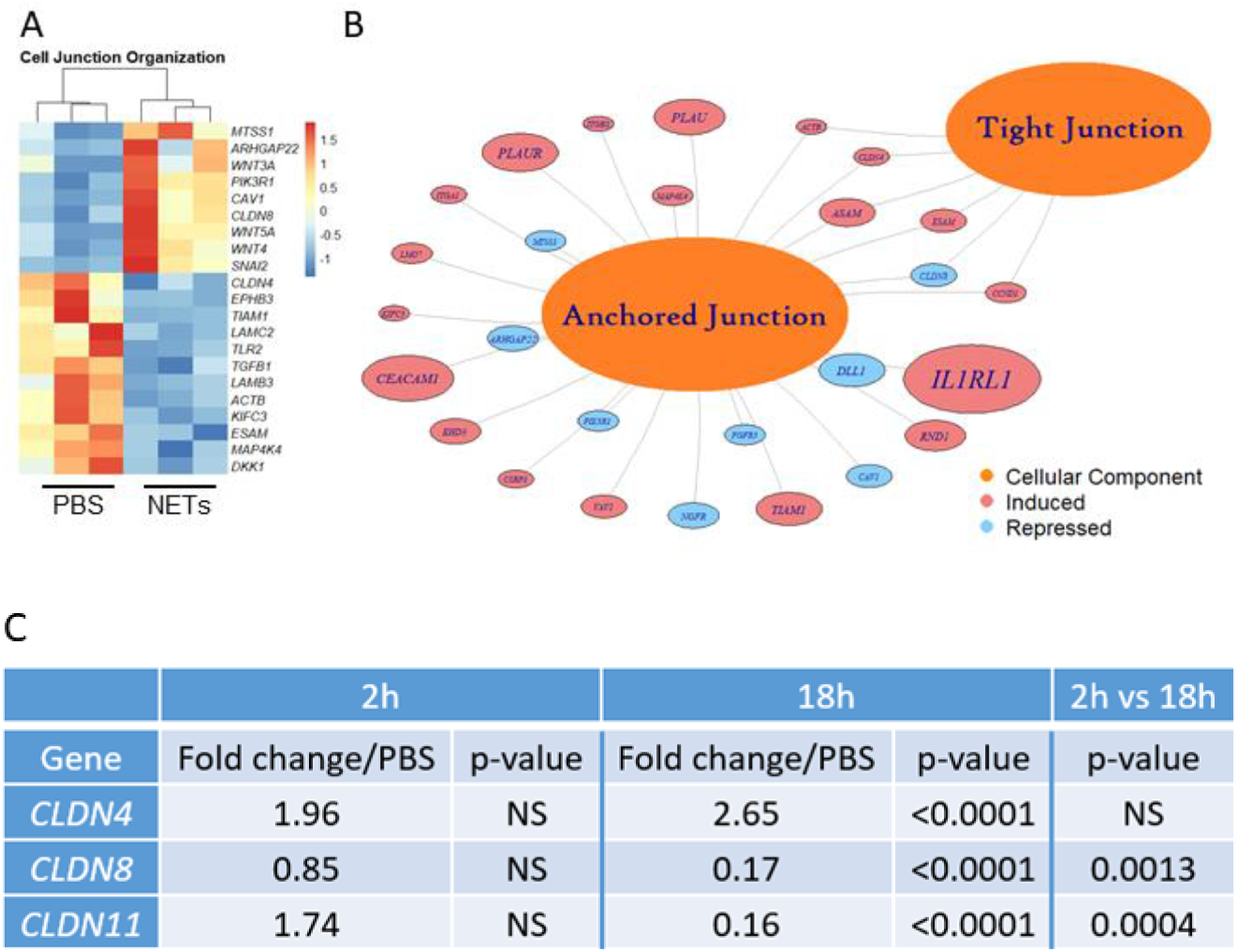
NETs alter epithelial cell junction genes. HBE grown at ALI exposed to PBS control or 5μg/ml NETs in the apical compartment in triplicate wells. **(A)** Differentially expressed genes involved in cell junction organization were visualized in a z-scored normalized heat map of fragments per Kilobase of transcript per million mapped reads (FPKM) demonstrate significant changes in HBE exposed to NETs for 18 hours. **(B)** Functionally enriched RNAs for cellular components of “anchored junctions” and “tight junctions” in HBE impacted by NET exposure for 18 hours, analyzed using Topp Gene (experiments were performed in triplicate HBE donors=3, NET donors=3). **(C)** Changes in junctional RNAs after 2 or 18 hour exposure to NETs were confirmed by RT-PCR and normalized to 18S. Data analyzed by unpaired t-test (experiments were performed in triplicate, HBE donors=4, NET donors=3).

Located in cell-cell tight junctions, claudins are tetraspan transmembrane proteins that regulate paracellular diffusion of ions, anions, and solutes (Schlingmann, Molina & Koval, 2015). Due to their importance in epithelial barrier function, we chose to focus on our RNA sequencing findings of changes in claudin gene expression. Claudin-4, a tight junction sealing protein and controller of anion flux, significantly increased with NET exposure after 18 hours (Van Itallie, Rahner & Anderson, 2001). In contrast, claudin-8 and claudin-11 were significantly downregulated by NETs at 18 hours (Figure 3C). Previously in the kidney, claudin-4 was shown to require claudin-8 for tight junction localization, where it forms an anion channel (Hou et al., 2010). Claudin-11 is expressed in renal, gastrointestinal and reproductive epithelia, and in the stria of the inner ear (Amasheh, Fromm & Günzel, 2011). Moreover, in Sertoli cells, claudin-11 localization within tight junction strands contributes to the formation of the blood-testis barrier (Morita et al., 1999). Disruption of barrier function and increased flux caused by NET exposure (Figure 1) is a key finding of our work. These results indicate NETs have selective and specific effects on junctional proteins in HBE.

To visualize the impact of NETs on specific bronchial epithelia junctional proteins, we performed confocal microscopy on wtCFBE41o- and HBE cells exposed to NETs for 18 hours, focusing on staining patterns of the critical and well-characterized E-cadherin protein (Figure 4). High-dose NET exposure led to the formation of gaps in the previously confluent wtCFBE41o-monolayer (Figure 4A-B). Images revealed partial loss of E-cadherin mediated cell-cell attachments in HBE monolayers exposed to NETs (Figure 4C-D). Empty space volume (ESV) calculated on these images was significantly increased in wtCFBE41o-exposed to NETs and trended towards increased in the HBE exposed to NETs, likely due to loss of cell-cell detachment with loss of E-cadherin (Supplemental Figure 3A-B). Disruption in nuclear organization in epithelia were evident in nuclear height maps in wtCFBE41o- and HBE exposed to NETs (Supplemental Figure 3C-D). Western blotting demonstrated significant reductions of E-cadherin protein in wtCFBE41o-(Figure 4E-F) and in HBE (Figure 4G-H) in response to NETs compared with control. Pretreatment of NETs with DA had no effect on loss of E-cadherin at 18 hours in wtCFBE41o-cells (Supplemental Figure 3A-B). NET-driven decreases in HBE E-cadherin were time-dependent with a trend toward reduction at 6 hours and a significant decrease in E-cadherin protein of 66.9% (p=0.0052) at 18 hours (Supplemental Figure 3C-D).

**Figure 4.**
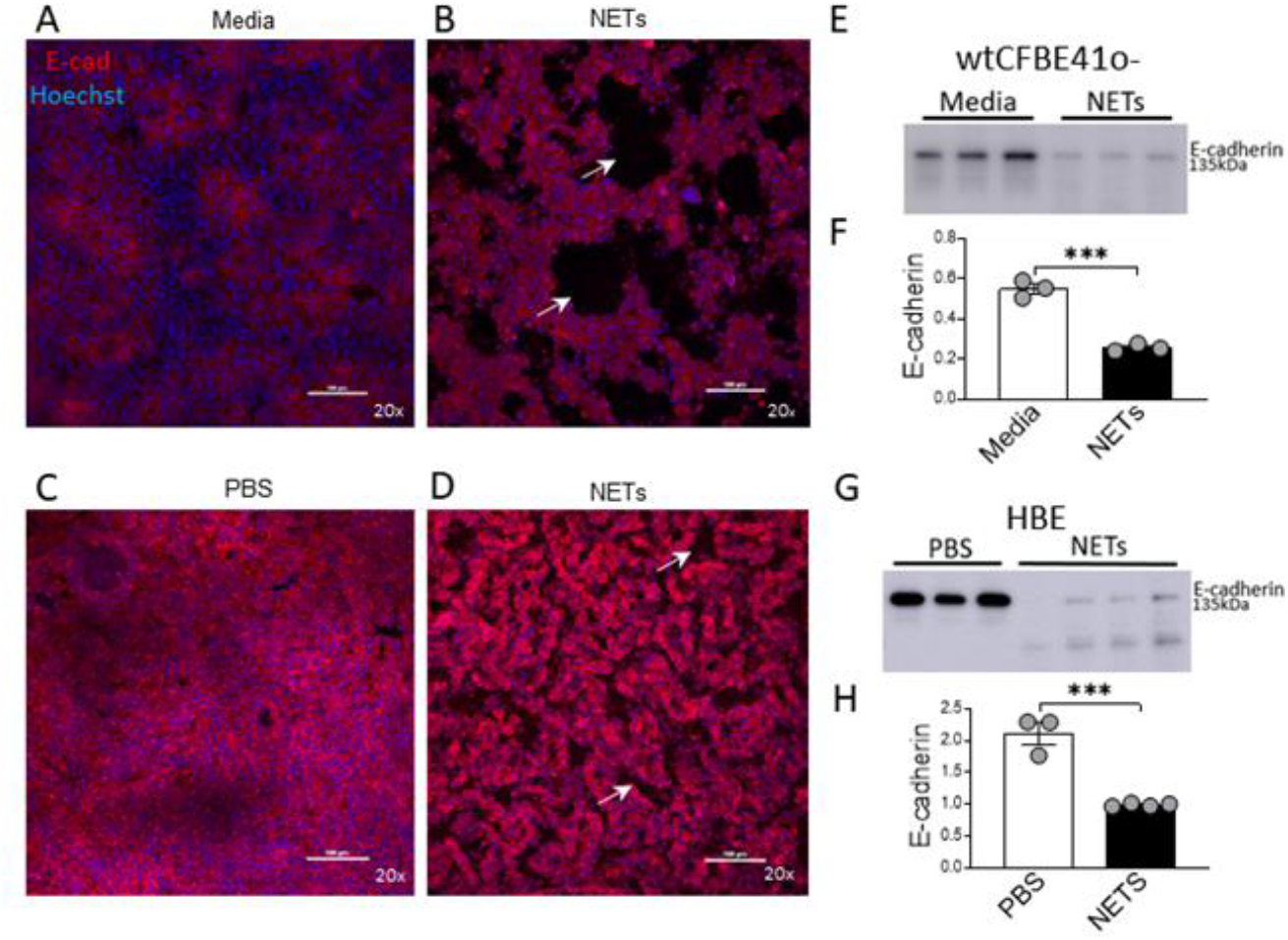
NETs disrupt epithelia and decrease E-cadherin. Confocal microscopy images (20x) of human epithelia stained for E-cadherin/AF647 and DNA/Hoechst. **(A)** wtCFBE41o-grown to confluence exposed to media or **(B)** 16μg/ml NETs for 18h, the latter with visible holes (arrows) in the monolayer. **(C)** HBE grown at ALI exposed to PBS or **(D)** 5μg/ml NETs for 18h, the latter with disruptions (arrows) in the monolayer. **(E)** Representative western blot and **(F)** corresponding analysis of E-cadherin protein from wtCFBE41o-lysates after 18h incubation with PBS or 16μg/ml NETs. E-cadherin normalized to C4-actin. Data analyzed by unpaired t-test (***p=0.0004, experiments performed in duplicate, NET donors=2). **(G)** Representative western blot and **(H)** analysis of E-cadherin protein from HBE lysates after 18h incubation with PBS or 5μg/ml NETs. E-cadherin normalized to C4-actin. Data analyzed by unpaired t-test (experiments were performed in triplicate, HBE donors=3, NET donors=3). ***p<0.001

### Alpha-1 antitrypsin limits NET-driven E-cadherin degradation

NETs are rich with serine protease enzymes, which exert antimicrobial effects (Urban et al., 2009). Given the complex structure of NETs and their associated proteases, we tested the susceptibility of NET proteases to inhibitors. Protease inhibitors decreased NET protease activity in a concentration-dependent manner in the absence of epithelial cells (Supplemental Figure 4A-C). NETs from three donors were separately exposed to A1AT, CG inhibitor (CGI), NE inhibitor sivelestat (SIV), pan-protease inhibitor III (PPI), or heat-inactivated NETs (HIN) for one hour. Heat reduced the activity of CG, PR3, and NE in NETs by 90-100%. A1AT reduced all three NET serine protease activities. CGI was a potent and specific inhibitor of CG, while SIV was most effective against NE and PR3. PPI was a less effective inhibitor overall. Reductions in activity differed across NET donors, suggesting varying susceptibility to protease inhibition. We also tested the effect of A1AT and NETs in the apical supernatant of HBE, where endogenous molecules may alter activity or compete for binding. A1AT was an effective NET serine protease inhibitor in the presence of HBE, significantly decreasing NE, CG, and PR3 activity (p<0.0001) in apical supernatants at 18 hours (Figure 5A-C).

**Figure 5.**
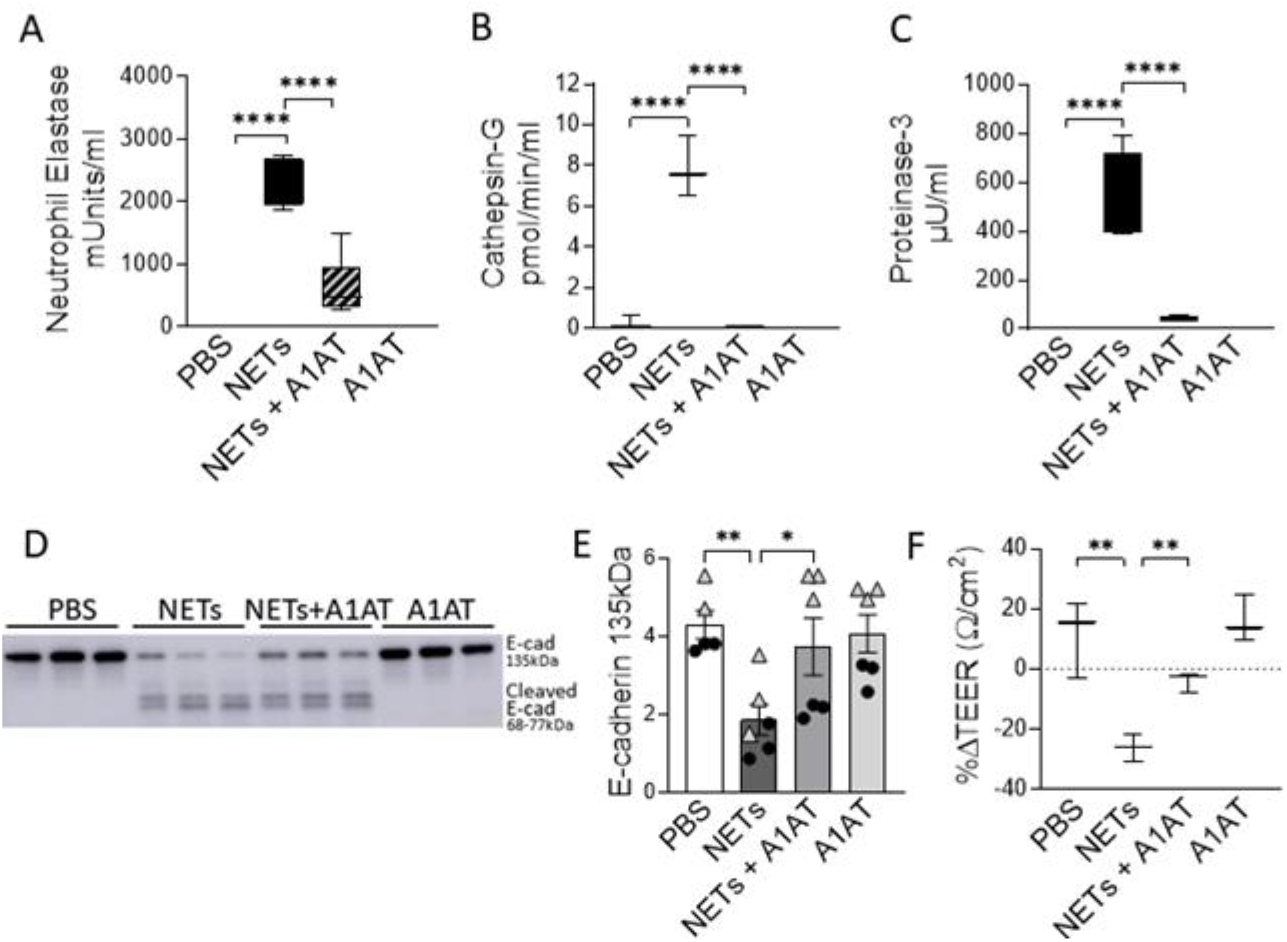
Alpha-1 antitrypsin reduces NET-driven degradation of E-cadherin. HBE were exposed to PBS, 5μg/ml NETs, NETs+100μg/ml A1AT or A1AT for 18h in triplicate wells. **(A-C)** Activity of NE, CG, and PR3 were reduced in the apical supernatants of HBE exposed to NETs + A1AT. Data analyzed by one-way ANOVA followed by Bonferroni’s multiple comparisons test (experiments=3, HBE donors=4, NET donor=2). (**D-E)** Representative western blot and analysis of E-cadherin protein from HBE lysates demonstrate that NET exposure significantly decreases (p=0.0016) the concentration of full-length E-cadherin (135kDa) and increases (p=<0.0001) cleaved E-cadherin (68 and 72kDa) compared to PBS control. A1AT significantly *limits* (p=0.0487) NET-driven reductions in full-length E-cadherin protein. Western blots normalized to C4-actin and analyzed by one-way ANOVA followed by Bonferroni’s multiple comparisons test (experiments=2, HBE donors=2, NET donors=3). **(F)** HBE exposed to NETs + A1AT had higher TEER than HBE exposed to NETs alone (representative graph). Data analyzed by unpaired t-test (experiments=3, HBE donors=3, NETs=3). *p<0.05, **p<0.01, ****p<0.0001.

To test the role of NET proteases in the degradation of E-cadherin, we exposed HBE to NETs with and without A1AT pretreatment. NETs caused cleavage of E-cadherin into 68, 72, and 77kDa fragments not observed in HBE exposed to PBS or A1AT alone. A1AT pretreatment of NETs significantly maintained full-length E-cadherin (135kDa) in HBE exposed to NETs (Figure 5D-E). A1AT decreased apoptosis in HBE exposed to NETs with significant decreases in full length caspase 9 and increased cleaved caspase 9 seen in Figure 5F-G. Next, we sought to determine if A1AT led to a functional stabilization of the electrical resistance of epithelial monolayers exposed to NETs. Importantly, pretreatment of NETs with A1AT preserved resistance compared to HBE exposed to NETs (Figure 5H). To visualize how NETs alter cell junctions on an ultrastructural level we performed transmission electron microscopy on HBE exposed to NETs for 18 hours in the presence and absence of A1AT. We found that NETs exposure caused gaps between cells at their tight junctions. A1AT limited NET-induced changes and preserved the close approximation between tight junctions in neighboring cells (Figure 6).

**Figure 6.**
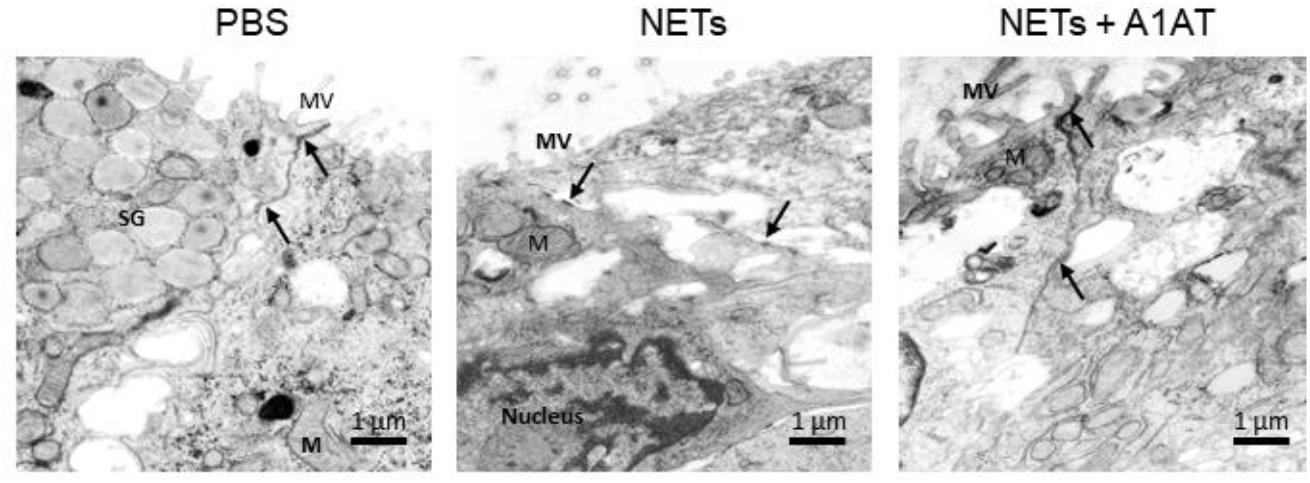
Alpha-1 antitrypsin preserves epithelial cell junction integrity disrupted by NETs. Representative images from transmission electron microscopy of HBE exposed to PBS, 5μg/ml NETs, NETs + A1AT or A1AT for 18 h. NET exposure resulted in partial disruption of epithelial tight junctions. A1AT helped maintain tight junctions in HBE exposed to NETs. (M=mitochondria, MV=microvilli, SG=secretory granules in a goblet cell, (→) =cell junction, experiments=1, HBE donors=1, NET donors=1).

### A1AT Complexes with NET Neutrophil Elastase

While A1AT binds and inactivates serine proteases from neutrophils, it is unclear whether A1AT binds NET proteases directly (Knoell et al., 1998). To demonstrate direct binding of A1AT to NET NE, we performed western blots on supernatants of HBE exposed to NETs with and without A1AT pretreatment. In blots probed for A1AT, 51 and 55kDa A1AT bands and an 80kDa A1AT:NE co-localizing band were detected (Figure 7A). A1AT alone was not detected by western blot in the supernatants of HBE exposed to NETs (Figure 7A). In blots probed for NE, a 29kDa NE band and the A1AT:NE complex were detected (Figure 7B). Most of the NE in NETs pretreated with A1AT co-localized with A1AT (Figure 7B). To further demonstrate NE:A1AT binding, we performed co-immunoprecipitation and detected the NE:A1AT complex (Supplemental Figure 4D).

**Figure 7.**
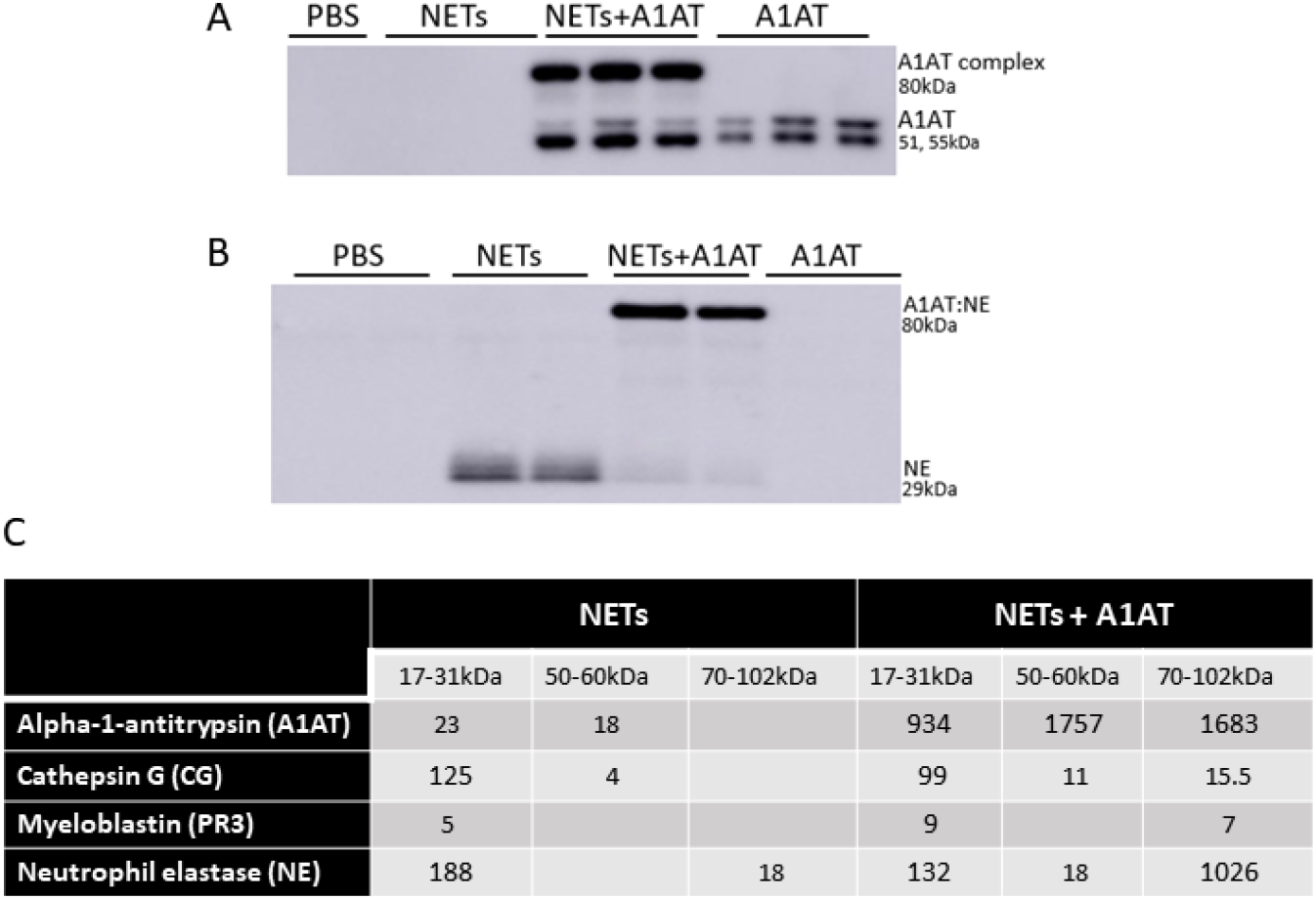
Alpha-1 antitrypsin binds serine proteases in NET exposed HBE. HBE grown at ALI were exposed to PBS, 5μg/ml NETs, NETs+100μg/ml A1AT or A1AT for 18h in triplicate wells. **(A)** Western blot of HBE apical supernatants probed with anti-A1AT antibody demonstrate the colocalization of A1AT (51-55kDa) with a 29kDa serine protease in NETs. **(B)** Western blot of HBE apical supernatants probed with NE antibody demonstrate the specific colocalization of A1AT with NE (29kDa) in NETs forming an 80kDa complex. **(C)** Mass spec. of bands shown on western blot in A & B confirm that HBE exposed to NETs have PSMs that correspond to CG, PR3, and NE at the expected 17-31kDa size. HBE exposed to NETs in the presence of A1AT demonstrate PSMs corresponding to NE and A1AT at 70-102kDa indicating formation of an A1AT:NE complex (experiments=3, HBE donors=3, NET donors=3).

To confirm our findings that NET NE complexes with A1AT we analyzed supernatants of HBE exposed to NETs with and without A1AT by mass spectrometry (Figure 7C). There were peptide spectral matches (PSMs) for CG, PR3, and NE detected at 17-31kD. There was a low level of full length A1AT detected, potentially from NETs or secreted by HBEs. There was no detectable A1AT:NE complex in the absence of exogenous A1AT (Figure 7C). There were PSMs for CG, PR3, NE and A1AT at 70-102kDa in HBE supernatants exposed to NETs pretreated with A1AT, which represent the individual proteases complexing with A1AT. Also in the HBE supernatant exposed to NETs pretreated with A1AT were peptide fragments of A1AT at 17-31kDa, likely the result of partial cleavage by other NET proteases. These mass spectrometric findings support the notion that A1AT directly forms complexes with NET proteases and limits NET-induced degradation of E-cadherin and disruptions in epithelial barrier function.

To determine if the source of endogenous A1AT was HBE, NETs or both we measured A1AT concentrations by ELISA in NETs alone and in the supernatants of HBE exposed to PBS or NETs. We found that NETs alone had consistent concentrations of A1AT protein across healthy donors (Supplemental Figure 5). A1AT was detected by ELISA in HBE exposed to both PBS and NETs, indicating both HBE and NETs were contributing endogenous A1AT. HBE exposed to NETs did not have additive levels of above that seen in HBE exposed to PBS (Supplemental Figure 5). We postulate that NE in NETs is cleaving A1AT in the supernatant. This is suggested by our mass spectrometry findings in Figure 7C in which we see PSMs for A1AT at band sizes of 17-31kDa, which is significantly smaller than the size of full length A1AT (51, 55kDa) and NE cleavage of A1AT is described in the literature (Janciauskiene et al., 2018).

## Discussion

NETs are associated with lung injury in many diseases, but the mechanisms of injury remain unclear (Lefrançais et al., 2018, Narasaraju et al., 2011). Our results demonstrate several pathologic changes induced by NET exposure.

NETs were sufficient to reduce electrical resistance in human airway cells and produce an increase in the paracellular flux of macromolecules. NET exposure activated transcription of genes promoting apoptotic cell death, which was confirmed by increased cleavage of the apoptosis initiator caspase-9 protein. In addition to NET-induced cytotoxicity, we observed selective changes in expression of tight junction genes including *CLDN4, CLDN8* and *CLDN11* in surviving epithelial cells. Bronchial epithelia exposed to NETs had visible disruptions in E-cadherin staining and reductions in E-cadherin protein in cell lysates. A1AT partially blocked NET-induced E-cadherin degradation, epithelial apoptosis and TEER reduction, emphasizing the role of NET serine proteases in epithelial injury. Collectively, these findings support the hypothesis that NETs induce epithelial cell death and regulate the expression of specific junctional proteins leading to disrupted epithelial barrier integrity (Figure 8).

**Figure 8.**
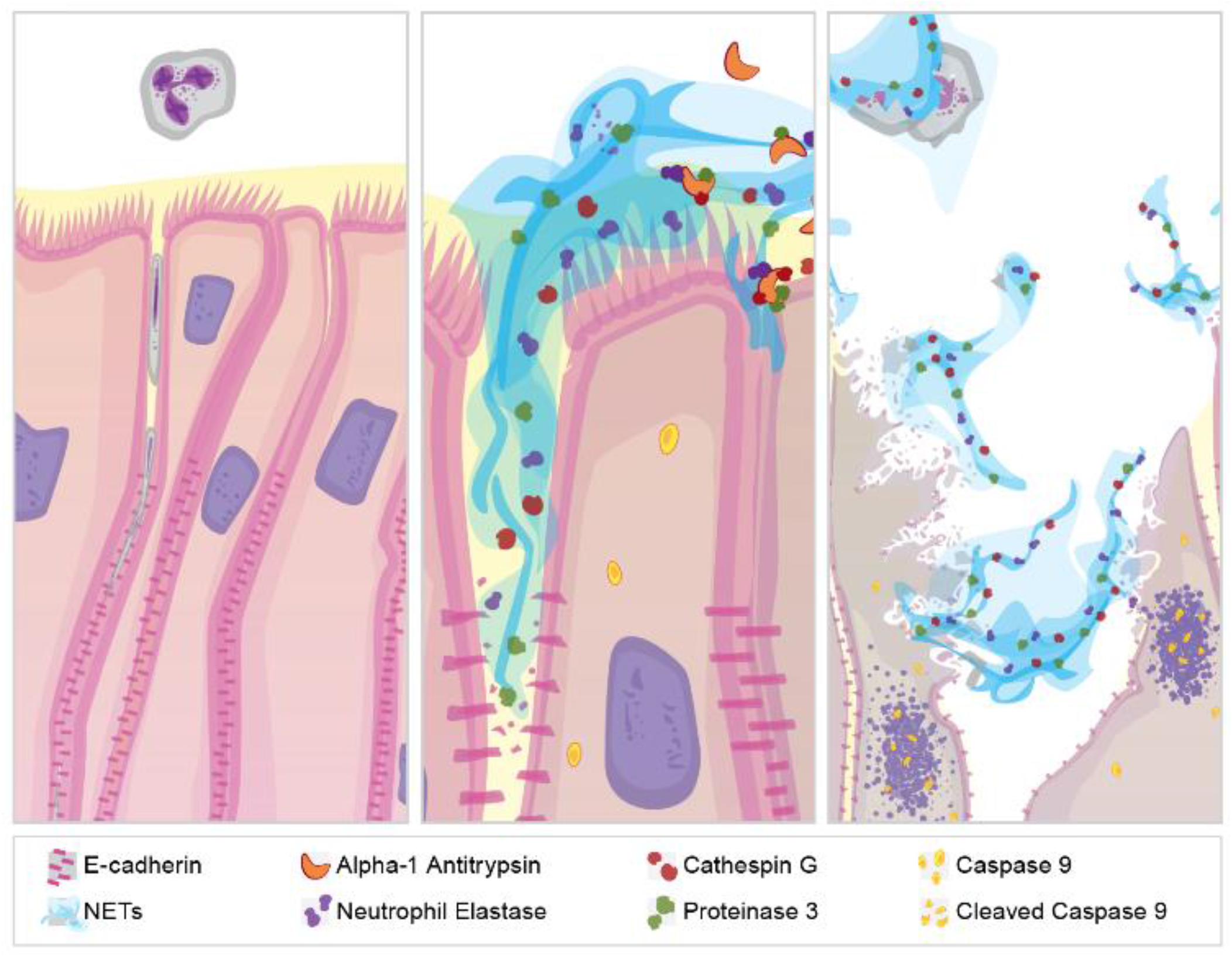
NET effect on airway epithelial barrier function. **(A)** Pseudostratified bronchial epithelia form a monolayer with cell-cell contact via junctional proteins. This epithelial barrier tightly controls the paracellular flux of molecules and cells. **(B)** Various stimuli cause neutrophils to expel NETs which contain biologically active proteases that degrade adherens junction protein E-cadherin creating gaps in the monolayer. A1AT limits NET degradation of E-cadherin to restore barrier function. C) NETs also disrupt barrier function by causing epithelial cell death via apoptosis.

Our findings have important clinical implications as elevated airway NET levels and disruption of epithelial barrier function have been observed in asthma and CF, but mechanistic links are lacking (Lachowicz-Scroggins et al., 2019, Xiao et al., 2011, Martínez-Alemán et al., 2017, Hudock KM, Imbrogno M, Collins MS, Brewington JJ, Kramer EL, O’Shaugnessy RA, Ostmann AJ, Clancy JP., 2020). Direct cleavage of junctional proteins or changes in expression of junctional molecules, both of which we demonstrate, have been reported to alter barrier function in lung diseases and infections (Reboud et al., 2017, Goto et al., 2000, Wray et al., 2009). Evidence for subtle barrier dysfunction in CF comes from glucose gradients in the blood that reflect airway concentrations, changes in claudin-3 expression in CF epithelia, and alterations in paracellular flux across CF epithelia (Molenda et al., 2014, Molina et al., 2017). Similar findings of barrier function disruption have been reported in asthma, including altered junctional protein expression, which is exacerbated by infection (Lachowicz-Scroggins et al., 2019). Although beyond the scope of this work, we postulate that changes in junctional RNAs could also be due to indirect effects of NETs on epithelia, e.g., cell stress or NET-induced changes in epithelial cytokines (Molenda et al., 2014, Hudock et al., 2020). The impact of barrier dysfunction in the human airway changes the susceptibility of the lung epithelium to infection, allergens or injurious stimuli.

Airway epithelia are relatively resistant to apoptosis, but when it occurs, barrier function is disrupted leading to increased pulmonary injury (Abdul-Hafez, Mohamed & Uhal, 2019, White, 2011). NETs were previously demonstrated to cause multicaspase activation in an alveolar adenocarcinoma cell line and, herein, we demonstrate that NETs induce apoptosis in primary human bronchial epithelia (Saffarzadeh et al., 2012). Individual NET components did not cause the cytotoxicity produced by NETs in airway epithelia despite isolated histones or neutrophil proteases producing cell death in other models (Saffarzadeh et al., 2012, Suzuki et al., 2005). These findings suggest that NET-induced cell death is likely multifactorial. Many of the pro-apoptotic transcriptional changes observed in our NET-exposed human bronchial epithelia included upregulation of proinflammatory cytokine pathways (Hudock et al., 2020). Future work will determine if NET-induced cytokine changes drive the airway epithelial cell apoptosis observed in our models (Shen et al., 2017, Hudock et al., 2020).

A key finding of this report is that NET exposure selectively cleaves E-cadherin, a junctional protein expressed at the basolateral surface of airway cells and an important regulator of monolayer integrity (Tian et al., 2011). The importance of E-cadherin in bronchial epithelia barrier function has been shown in the 16HBE41o-cell line in which blockade of E-cadherin decreased electrical resistance of the epithelial monolayer and enhanced susceptibility to adenoviral infection (Man et al., 2000). E-cadherin has multiple potential cleavage sites for NET serine proteases. Although purified NE, CG, and PR3 have been shown to cleave E-cadherin in bronchial cell lines *in vitro*, to our knowledge, this is the first demonstration that human NETs cleave E-cadherin resulting in altered barrier function in primary human bronchial epithelia (Boxio et al., 2016, Evans et al., 2002). The ramifications for NET-induced decreases in full length E-cadherin in the human bronchial epithelia are profound and could uncover a novel role for NETs in driving epithelial-mesenchymal transition (by increasing β-catenin), facilitating invasion of pathogens (similar to sheddases) and increasing metastases in cancer (due to loss of cell-cell adhesion) (Nawijn et al., 2011, Brune et al., 2015, Devaux, Mezouar & Mege, 2019, Bruner, Derksen, 2018).

Another critical result of this study is demonstration of novel therapeutic effects of A1AT. Protease-antiprotease imbalance has been postulated to contribute to many pulmonary diseases, including COPD and alpha-1 antitrypsin deficiency (Chillappagari et al., 2015, Fazleen, Wilkinson, 2021). Increased neutrophil numbers and NE activity are associated with severe asthma and bronchodilator resistance (Crisford et al., 2021, Ray, Kolls, 2017). NE levels and activity are elevated in bronchoalveolar lavage fluid (BALF) from infants with CF, even in the absence of infection, and are associated with early bronchiectasis (Khan et al., 1995, Sly et al., 2013). Small clinical trials of inhaled A1AT in CF demonstrated some benefits including decreases in Pseudomonas aeruginosa, proinflammatory cytokines (e.g., IL-8), neutrophils and/or neutrophil markers (e.g., NE). Notably, these studies did not show any significant increase in lung function, but study drug was given for short intervals (1-4 weeks), and a more realistic outcome may have been to limit progressive lung function decline. (Brennan, 2007; Griese et al., 2007; Martin et al., 2006; McElvaney, N. G. et al., 1991; McElvaney, Noel G., 2016) Whether NET-derived NE (and/or other proteases) contribute to these relationships between NE and airway disease pathology is unknown. We examined whether the naturally occurring serine protease inhibitor A1AT was sufficient to impact NET damage to airway epithelia integrity. Colocalization of fluorescently labeled A1AT and NE in NETs, generated *ex vivo* from alpha-1 antitrypsin deficient patients receiving recombinant A1AT therapy, has been reported but the functional implications have not (Frenzel et al., 2012). In addition to demonstrating that A1AT binds NET serine proteases, we show that it inhibits NET protease activities. The importance of our finding that NET NE complexes with A1AT is that NE is known to cleave A1AT, which causes a change in A1AT conformation. This produces formation of a covalent bond between NE and A1AT that renders both molecules inactive (Huntington, Read & Carrell, 2000, Brantly, 2002, Janciauskiene et al., 2018). Furthermore, this could lead to NET depletion of endogenous A1AT further suggesting that A1AT augmentation therapy may be critical to limit destruction of lung tissue in conditions with high NET burden. Moreover, we demonstrate that A1AT impacts other NET effects on bronchial epithelial including limiting apoptosis. This may occur via A1AT inhibition of caspase 3 as reported in lung endothelium (Petrache et al., 2006). Our findings show potential for an anti-protease therapy to disrupt NET effects, offering new therapeutic directions to ameliorate diseases with a high NET burden.

A strength of our study was that we used NETs from eight donors and primary airway epithelia from 10 human donors reflecting the broad variability between individual subjects. Previously, we and others have reported variability in the concentration and protease activities of NETs from different human subjects, which contributes to inconsistency in results between groups of investigators (Hudock et al., 2020, Hoffmann et al., 2016). The evidence we present of similar epithelial responses to NETs across both primary airway epithelial and a bronchial epithelial cell line reinforces the validity of our findings and likely relevance to *in vivo* conditions. Notably, our most critical findings of NET cleavage of E-cadherin and disruption of epithelial barrier function with amelioration by A1AT were consistent across NET and epithelial donors. An additional strength is that our demonstration of a cohesive sequence of NET-induced events involving epithelial disruption, apoptosis, and destruction of the monolayer has implications in both acute and chronic lung diseases, e.g., ARDS and CF. Our work offers initial proof that NETs, with their complex structure, are more injurious than the sum of their parts.

Limitations of our studies include defining NET “dosage” by DNA content alone, which despite acceptance in the field, does not account for 100+ other NET components (Saffarzadeh et al., 2012, Chapman et al., 2019). We have attempted to address this limitation by using multiple primary HBE cell and NET donors. Additionally, our experiments do not factor in the positive effects of NETs, which include limiting the spread of pathogens (McDonald et al., 2012, Thanabalasuriar et al., 2019). Although our *in vitro* studies are consistent across multiple airway epithelial cell and NET donors, further studies will be needed to verify if NETs cause epithelial cell cytotoxicity *in vivo*. We do not account for the potential differences in our NETs generated from peripheral blood neutrophils compared to NETs generated from neutrophils within tissues. This is particularly relevant in the lung in which transmigration and the microenvironment are known to alter transcription, translation and key functions, e.g, bacterial killing, of the neutrophil (Forrest et al., 2018; Margaroli et al., 2021).

In summary, we present a novel paradigm in which NETs are sufficient to disrupt human airway epithelial barrier function by causing altered regulation of junctional protein expression, cleavage of E-cadherin and bronchial epithelial apoptosis, the latter two of which are limited by exogenous A1AT. NET-driven injury to the airway epithelium is a pathway of inflammatory damage relevant to many airway diseases and further investigations may lead to novel therapeutic strategies to mitigate lung function decline in CF and asthma. Finally, NET-driven decreases in E-cadherin have potential consequences that extend beyond disruption of epithelial barrier function and future studies will delve into this biology.

## Materials and Methods

RESOURCES AVAILABILITY Lead Contact for requests concerning resources is Kristin Hudock

### Generation of human NETs

Polymorphonuclear (PMN) cells were isolated from the peripheral blood of healthy adult human subjects via negative bead selection using the MACSxpress Neutrophil Isolation Kit, Human #130-104-434 Miltenyi Biotec (RRID:SCR_008984) as previously described (Hudock et al., 2020) and according to manufacturer’s direction. In brief, for each NET isolation, 60ml blood was collected into 10ml EDTA (K2) tubes (BD Bioscience #366643). NETs from six different donors were utilized to capture the biologic variability contributed by individuals’ neutrophils. Whole blood was incubated with negative selection magnetic beads, and neutrophils were isolated using magnetic separators (Miltenyi #130-101-319). Purified PMNs were plated in 100mm cell culture dishes, 7.5×10^7^/15ml RPMI+3%FBS and stimulated with 25nM phorbol 12-myristate 13-acetate (PMA; Millipore-Sigma #P8139) Plates were incubated at 37°C, 5% CO_2_ for 4 hours to induce NET generation (Reinhold et al., 2010). Following incubation, the viscous surface layer of NETs was washed with PBS for HBE studies, or media for wtCFBE41o-experiments, scraped from the dish, and centrifuged 1,200rpm 10m, to yield cell-free human NETs (Hudock et al., 2020). NET dosage was defined by DNA concentration using QuantiFluor ONE dsDNA system Promega #E4871 and SYTOX green nucleic acid stain Invitrogen S7020 as previously described (Saffarzadeh et al., 2012, Hudock et al., 2020).

### Primary HBE and wild-type CFTR CFBE41o-cell culture and treatment

The CCHMC CF Research Development Program Translational and Model Systems Cores maintained, and provided wild-type CFTR (wtCFBE41o-) cells as previously described (Brewington et al., 2018). wtCFBE41o-cells stably transduced with wild-type CFTR were initially generated and described by Gruenert and colleagues, and were initially a kind gift from the University of Alabama at Birmingham (Haws et al., 1992). HBE from ten normal healthy donors were obtained from University of North Carolina Airway Cell Core using previously described methods (Fulcher et al., 2005). HBE at passages 2-3 were differentiated and grown at ALI on Corning® 3470 Transwell® polyester membrane cell culture inserts, 6.5 mm diameter, 0.4μm pore size, by the CCHMC Pulmonary Core as described (Brewington et al., 2018). The CCHMC, Pulmonary Biorepository Core, and University of Cincinnati IRBs approved primary HBE use. Epithelial cells were incubated with NETs from unrelated human donors, at doses of ∼5μg/ml, 18h, 37°C, 5% CO_2_. Higher dose NETs or other incubation periods were used in experiments where indicated. Dornase Alpha (DA), obtained from the CCHMC research pharmacy, was used at a final concentration of 0.5μg/ml to digest the NET DNA structure.

### Transepithelial electrical resistance (TEER)

TEER was measured across each epithelial cell layer in duplicate before and after exposure to NETs, controls, and other reagents using an Epithelial Volt/Ohm Meter EVOM2 and chopstick electrode set STX3, World Precision Instruments (RRID:SCR_008593) as previously described (Brewington et al., 2018). Results are expressed as percent change from baseline values.

### Collection and characterization of cell supernatants

Cells were exposed on their apical surface to controls, NETs or other test compounds in a final volume of 200μl per 6.5mm insert. Supernatants were collected from the apical surface, and aliquots were stored at -80°C. Assays were performed to measure human NE (Cayman Chemical 600610), CG (Cayman Chemical 14993, ABCAM 126780), PR3 (Cayman Chemical 9002021) (Zaidman et al., 2017), LDH (Promega). NETs contain LDH, which was subtracted from the total to determine the contribution by HBE. The NET LDH was measured by incubating NETs alone for 18h at 37°C. Across multiple experiments the mean was 675mU/ml. Human A1AT was measured by ELISA (R & D DY1268). All assays were performed per manufacturer’s instructions. Absorbance and fluorescence quantitation was performed using Molecular Devices FlexStation 3.

### Protein Isolation and Western Blot

Epithelial cells were lysed using cOmplete Lysis-M, (Roche Diagnostics #04719956001) and protein was harvested as described (Clancy et al., 2018). Protein concentrations were measured using DC Protein assay (BioRad #500-0114) with bovine serum albumin (BSA, Thermo-Fisher #BP7906) standards. Proteins were separated and immobilized using NOVEX electrophoresis and transfer apparatus, 4-20% Tris-glycine gels (Invitrogen #XPO420), and 0.45μm PVDF membrane (Thermo-Fisher #88518). Western blot was performed as previously described (Kramer et al., 2020). Protein lysates of 16HBE cells exposed for 2h to 5μM staurosporine, (Sigma-Aldrich #S6942) were used as a positive control for apoptosis. Antibodies for western blot were obtained from Cell Signaling Technology (CS) and Abcam (AB). We used Caspase-9 CS9502, E-cadherin CS3195, A1AT AB207303, NE AB68672, RIP1 CS3493, MLKL CS14993, p-MLKL CS91689 and C4 anti-actin (Seven Hills Bioreagents LMAB-C4). Secondary antibodies were rbHRP (CS7074) and msHRP (CS7076). Blots were developed using Cytiva ECL Prime detection reagent (Amersham #RPN2232). Images and analysis were performed using Amersham Imager 600.

### RNA isolation and sequence analysis

RNA sequencing analyses (RNA-seq) was performed on primary HBE data previously generated and described (Hudock et al., 2020). Differentially expressed genes had a fold change >2.5, a p-value (negative binomial Test) <0.01, and reads per kilobase million (RPKMs) >2 for 50% of the samples in at least one condition. Ingenuity Pathway Analyses (IPA) was used to predict activation or repression of known biological process or pathways and was used to visualize predicted “apoptosis” activation (Krämer et al., 2014). Functional Enrichment analyses was done using Toppfun to determine biological functions or cellular components enriched in genes differentially regulated following NETs exposure (Chen, J. et al., 2009). Significant, relevant biological functions or cellular components were visualized in a z-score normalized Fragments Per Kilobase of transcript per Million mapped reads (FPKM) heatmap using pheatmap (https://cran.r-project.org/web/packages/pheatmap/index.html) or a Kamada-Kawai layout generated network using igraph (https://cran.r-project.org/web/packages/igraph/index.html) (Csardi), respectively. Quantitative PCR was performed with TaqMan master mix and primers CLDN4 (Hs00976831), CLDN8 (Hs 04186769) CLDN11 (Hs00194440) normalized to EUK 18S rRNA AB (1207030) (Millipore-Fisher Scientific).

### Immunofluorescence microscopy

Cell culture monolayers were prepared as previously described (Clancy et al., 2018). Briefly, cells were fixed in 4% paraformaldehyde (Fisher Scientific #50-980-487), washed and incubated with mouse anti-human E-cadherin antibody (Cell Signaling Technologies), Alexafluor AF647 (Cell Signaling Technologies), and Hoechst 33342 nucleic acid stain (Invitrogen #H3570). Confocal Images were captured on a Nikon A1R GaAsP inverted SP microscope, and analyzed using IMARIS software.

### Transmission Electron Microscopy

Cultured cells growing on Corning Transwell 3470 filter membrane inserts were fixed *in situ* with 1 part cell culture media and 1 part 4% paraformaldehyde (PF), 4% glutaraldehyde (GA) and 0.2% CaCl_2_ in 0.1M sodium cacodylate buffer (SCB), pH 7.2 at RT for 10m, followed by fixation with 2% PF, 2% GA, and 0.1% CaCl_2_ in 0.1 SCB, pH 7.2, on ice for 30m. Samples were stored in fresh fixative at 4°C for up to 2 days. Fixed membranes were cut into 1-2 mm stripes and processed for electron microscopy as previously described (Yang et al., 2017, He et al., 2021). Electron micrographs of cultured cells were acquired using a Hitachi H-7650 TEM and an AMT CCD camera.

### Mass spectrometry

Modified proteomic methods previously established by Ziady et.al. were used (Chen, X. et al., 2011, Ziady, Kinter, 2009, Chen, J. et al., 2008). The database included A1AT, CG, PR3 (myeloblastin), and NE. The following conditions were tested: NETs and NETs + A1AT. Proteins from each condition were separated on a 4-20% Criterion™ Tris-HCL gel, and bands of interest were excised from the gel at three molecular weight ranges: 17-31kDa, 50-60kDa, and 70-102kDa. The samples were then subjected to in-gel trypsin digest (Trypsin Gold, Mass Spectrometry Grade. Promega. 20μg/ml). Samples were loaded in an UltiMate 3000 HPLC autosampler system (Thermo Fisher) and were eluted using reverse-phase column chromatography into an LTQ Velos Pro with a nanospray ion source. The mass analyzer used data-dependent settings with dynamic exclusion enabled: repeat count = 2; repeat duration = 10 seconds; exclusion list size = 100; exclusion duration = 10 seconds; exclusion mass width = 1.5 amu. Collision-induced dissociation (CID) was used to fragment peptides, and CID spectra were searched against a human FASTA database using the Proteome Discoverer™ software version 2.4. We used a 5 score cutoff in addition to a 1.0 XCorr threshold. The protein score is the sum of all peptide XCorrs for that protein above a certain threshold. This threshold is determined by the peptide charge and a set peptide relevance factor of 0.0 – 0.8. Proteins were quantified based on peptide spectral match (PSM). Each protein must have a minimum of 2 peptides.

### Immunoprecipitation

Immunoprecipitation of A1AT and NE proteins was carried out using supernatants from HBE incubated with PBS, NETs, 100μg/ml A1AT, and NETs pretreated with A1AT. Protein A Dynabeads (Invitrogen 10-001-D) were washed with PBS. Samples were incubated with washed beads 1h, 4°C. Pre-cleared samples were incubated with A1AT or NE antibodies (Abcam 207303, 68672) 1h, 4°C. Washed beads were added and incubated overnight at 4°C with rotation. Beads were washed twice in PBS, and proteins eluted with 1X Laemmli buffer (Boston Bioproducts #BP-111R) 100°C 10 minutes. Resulting proteins were run on an SDS PAGE gel (Invitrogen #XPO420), transferred to PVDF membrane (Thermo-Fisher #88518), and probed with corresponding NE or A1AT antibodies.

### Dextran flux measurements

wtCFBE41o-monolayers were grown to resistances of >600Ωcm^2^ on cell culture inserts. FITC-labeled dextran (molecular mass 4,000Da, Sigma-Aldrich #46944, 2.0mg/ml) in media +/-NETs was added to the apical compartment. At each time point, 100μl samples of the basolateral solutions were transferred to opaque 96-well plates. Fluorescence intensity (485nm excitation/530nm emission wavelengths) was measured using Molecular Devices FlexStation 3 plate reader. To determine dextran concentration, a standard curve was generated using FITC-dextran dilutions of 0.61-620μg/ml and unidirectional fluxes at each time point were calculated (Zaidman et al., 2017).

### Protease inhibitors and Individual NET Components

NETs were pre-incubated with A1AT protein from human plasma (Cayman Chemical #24560) 1h before exposure to HBE. Individual components of NETs were incubated with wtCFBE41o-to assess the effect of each on cytotoxicity and monolayer integrity. These included 5μg/ml human gDNA (Promega), 500ng/ml citrullinated histones (Cayman Chemical #501620), 100mU/ml human NE protein (Athens Research & Technology #16-14-051200, ART), 100μU/ml human PR3 protein (ART #16-14-161820) and 4.5mU/ml human CG protein (ART #16-14-030107). These were compared to media control, and 5μg/ml NETs. The concentration of individual NET components was chosen to reflect that found in our isolated human NETs. In epithelial cell-free experiments, NETs were pre-incubated with protease inhibitors: human A1AT (Cayman Chemical 24560), sivelestat (Cayman Chemical #177749), and cathepsin G inhibitor I (Cayman Chemical #14928) all at 1μg/ml and 100μg/ml, and protease inhibitor cocktail III at 0.1μM and 100μM (Alfa Aesar #J64283) (1h at 37°C) and assayed for protease activity (Korkmaz et al., 2008).

### Statistical Analysis

Statistical analyses were performed using GraphPad Prism software v7.04 and SAS v9.4. Groups were compared using one-way ANOVA with Bonferroni’s correction for multiple comparisons or a Student’s t-test where indicated. Data are expressed as means +/-SEM. A p-value of <0.05 was considered significant. Graphs include data from at least three separate experiments unless indicated. Each experiment included conditions run on triplicate or more wells (as noted in figure legends). As per GraphPad analysis, p values ≥0.05 are not significant, p-values <0.05 are denoted by *, p-values ≤0.01 are denoted by **, p-values ≤0.001 are denoted by ***, and p-values ≤0.0001 are denoted by ****. In each figure, the number of experimental repeats with similar results is noted. “NETs =“ refers to the number of normal human neutrophil donors from which NETs were isolated. “HBE donors =“ refers to donors of primary bronchial epithelia samples obtained from the University of North Carolina Airway Cell Core. A total of 10 HBE donors and 6 NETs donors were used in this study. A linear mixed effect model was used to evaluate whether wtCFBE41o-cells exposed to NETs had significantly different dextran flux over the course of 0-24h, compared to media control alone. If a difference existed, post-hoc comparisons were conducted to compare dextran flux between NETs and media control only group at each time point.

## Acknowledgements

Graphical abstract and illustrations by Laura Collins, Freiburg, Germany. Dr. Matt Kofron and the Confocal Imaging Core at Cincinnati Children’s for their help with imaging and analysis.

## Author Contributions

Designing research studies: KMH, MSC, MI; conducting experiments: MSC, MI, ES, MES, JM, HM, AJO, CLN; analyzing data: KMH, MSC, MI, ES, AZ, CLN, CRD, NZ, JS, YX; intellectual content; KMH, MSC, MI, ELK, JJB, AZ, SMO, EK, KG, CLN, NM, BCT, JPC; writing the manuscript: KMH, MSC, MI; revisions: all authors

## Study Approval

The study protocol, no. 2016-3837, was approved by the University of Cincinnati IRB. Healthy adult human subjects provided written informed consent prior to participation in our study.

