## Supplemental figures for "Alpha-1 Antitrypsin Limits Neutrophil Extracellular Trap Disruption of Airway Epithelial Barrier Function"

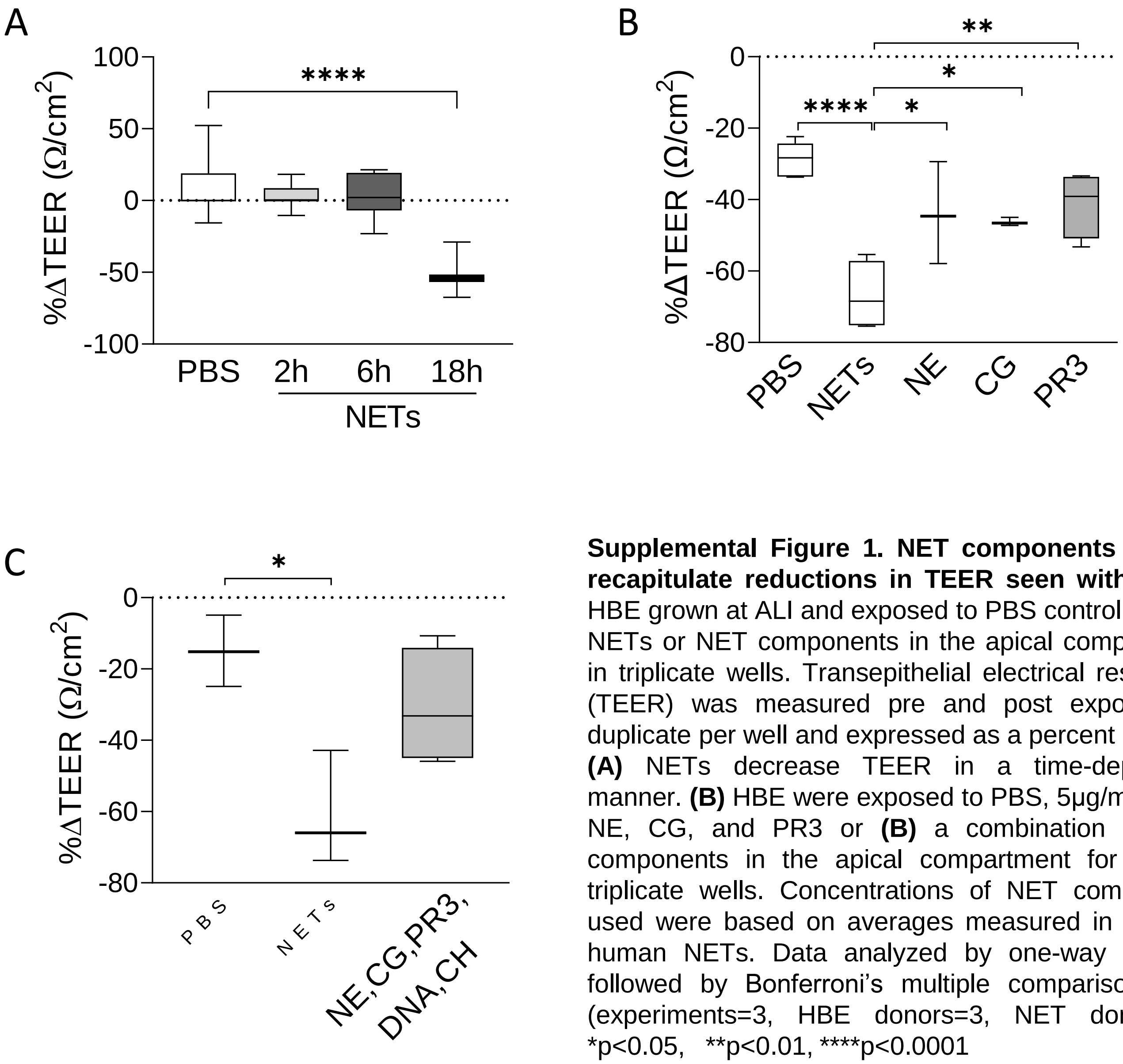


**Supplemental Figure 1. NET components do not recapitulate reductions in TEER seen with NETs.** HBE grown at ALI and exposed to PBS control, 5µg/ml NETs or NET components in the apical compartment in triplicate wells. Transepithelial electrical resistance (TEER) was measured pre and post exposure in duplicate per well and expressed as a percent change. **(A)** NETs decrease TEER in a time-dependent manner. **(B)** HBE were exposed to PBS, 5µg/ml NETs, NE, CG, and PR3 or **(B)** a combination of NET components in the apical compartment for 18h in triplicate wells. Concentrations of NET components used were based on averages measured in isolated human NETs. Data analyzed by one-way ANOVA followed by Bonferroni's multiple comparisons test (experiments=3, HBE donors=3, NET donors=4). *p<0.05, **p<0.01, ****p<0.0001

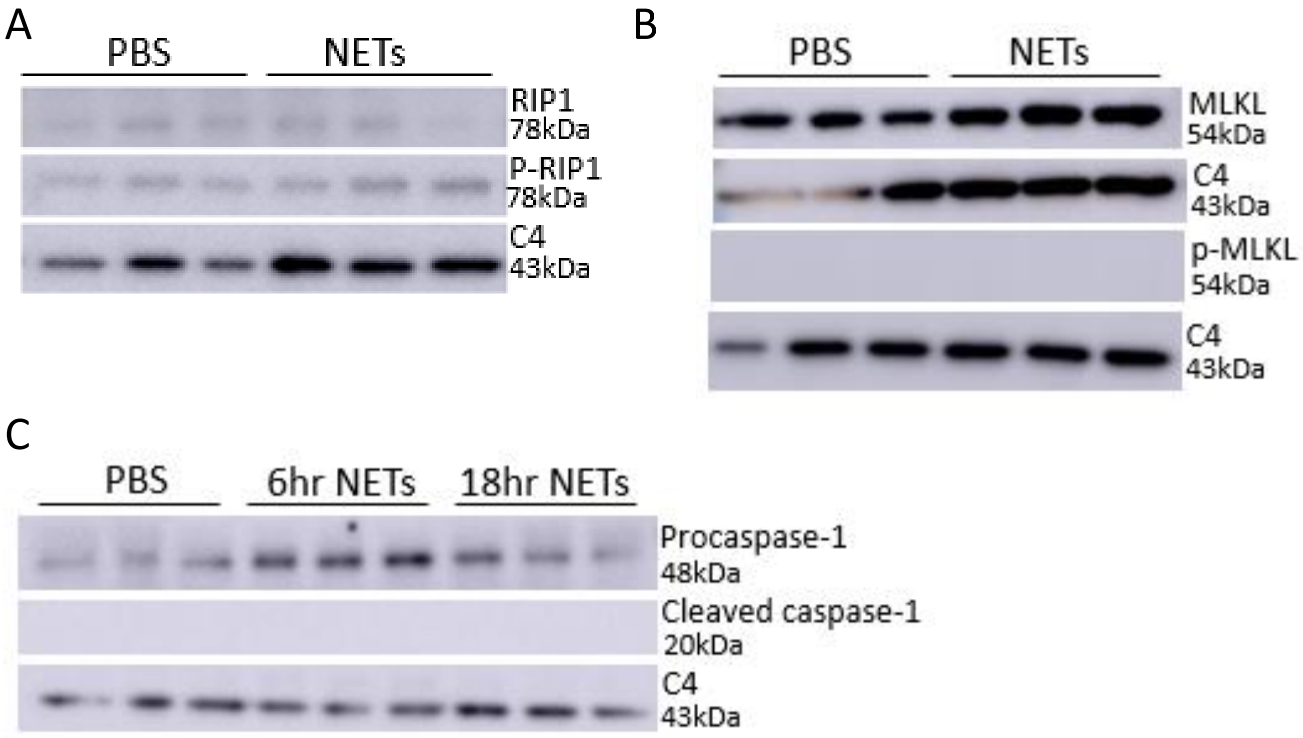


**Supplemental Figure 2. NETs do not induce necroptosis or pyroptosis in human bronchial epithelia.** HBE grown at ALI were exposed to PBS control or 5µg/ml NETs in the apical compartment for 18h in triplicate wells. **(A)** Representative western blot of receptor-interacting serine/threonine kinase 1 (RIP1), phosphorylated RIP1, **(B)** mixed lineage kinase domain-like (MLKL) and phosphorylated MLKL proteins of lysates from HBE after 18h NET exposure demonstrate no change. **(C)** Representative western blot demonstrates NET exposure for 6 or 18 hours do not cleave caspase-1 (20kDa) to activate pyroptosis in HBE. (experiments=2, HBE donors=2, NETs=3)

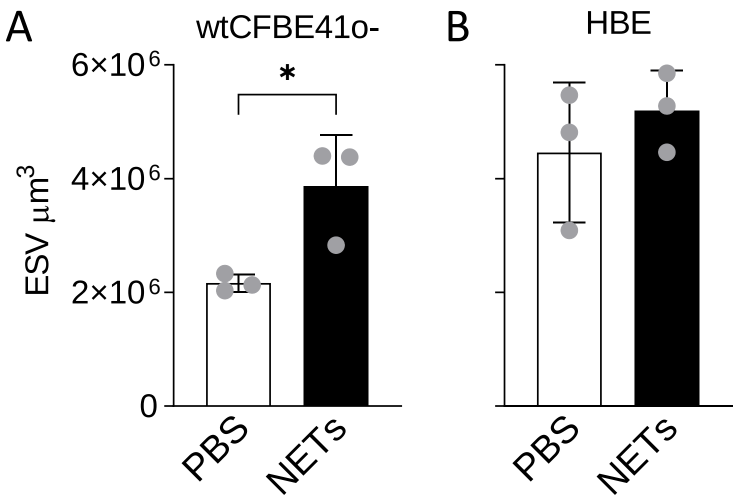


**Supplemental Figure 3. NETs Increase Empty Space Volume and Disrupt Nuclear Organization in Human Bronchial Epithelia.** wtCFBE41o- grown to confluence were exposed to media or 15µg/ml NETs in the apical compartment for 18h in triplicate wells. Primary normal human epithelia (HBE) grown at air-liquid interface (ALI) were exposed to PBS or 5µg/ml NETs in the apical compartment for 18h in triplicate wells. **(A-B)** Empty space volume was calculated using Imaris cell imaging software and represents disruptions in the epithelial monolayers. **(C-D)** Nuclear organization was assessed by mapping nuclear heights in Z-stacks.

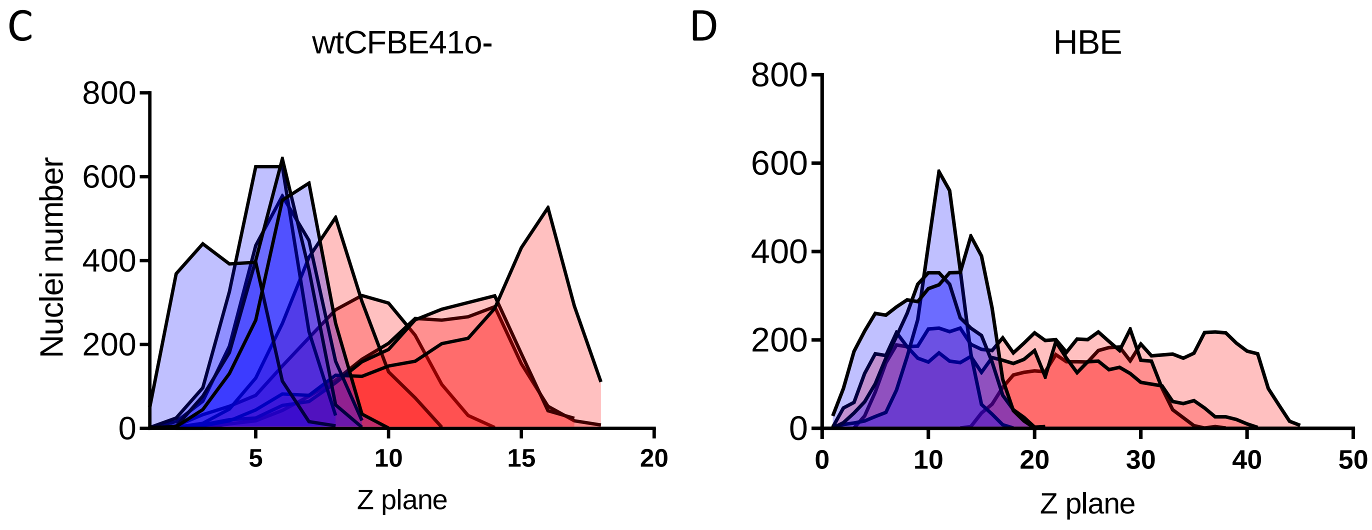

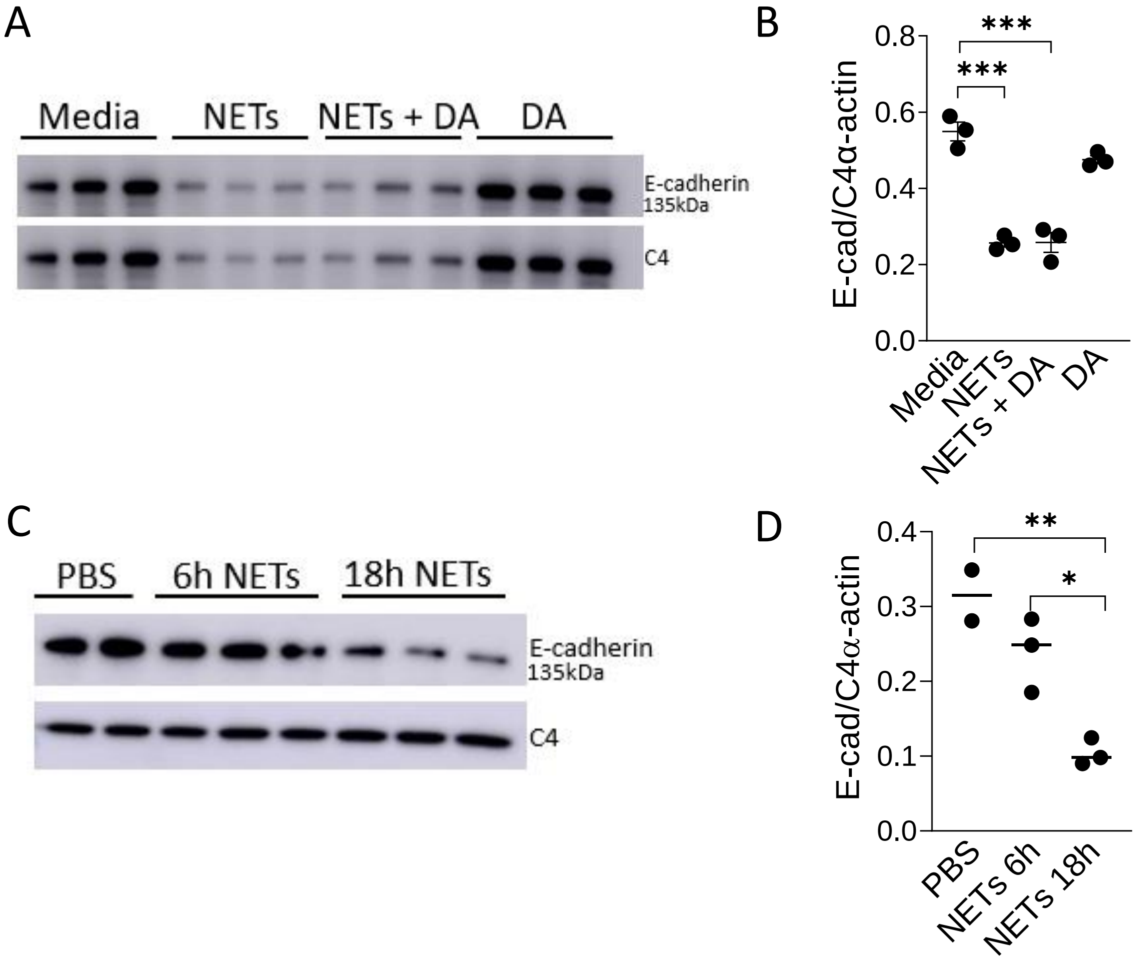


**Supplemental Figure 4. Dornase alpha does not alter NET driven reductions in E-cadherin.** wtCFBE41o- or HBE were exposed to media or PBS control (respectively), 5µg/ml NETs, NETs+0.5µg/ml DA, or DA in the apical compartment for 18h in triplicate wells. **(A-B)** Representative western blot with corresponding analysis of E-cadherin protein from lysates of wtCFBE41o- demonstrate that NETs reduced E-cadherin in the presence and absence of DA. **(C-D)** Representative western blot and corresponding analysis of E-cadherin protein from lysates of HBE exposed to NETs demonstrate a trend towards a decrease in full-length E-cadherin at 6h and a significant decrease at 18h, indicating E-cadherin is decreased by NETs in a time-dependent manner. E-cadherin was normalized to C4-actin. Data analyzed by unpaired t-test (experiment=1-3, HBE donor=1, NET donor=4) *p<0.05, **p<0.01, ***p<0.001.

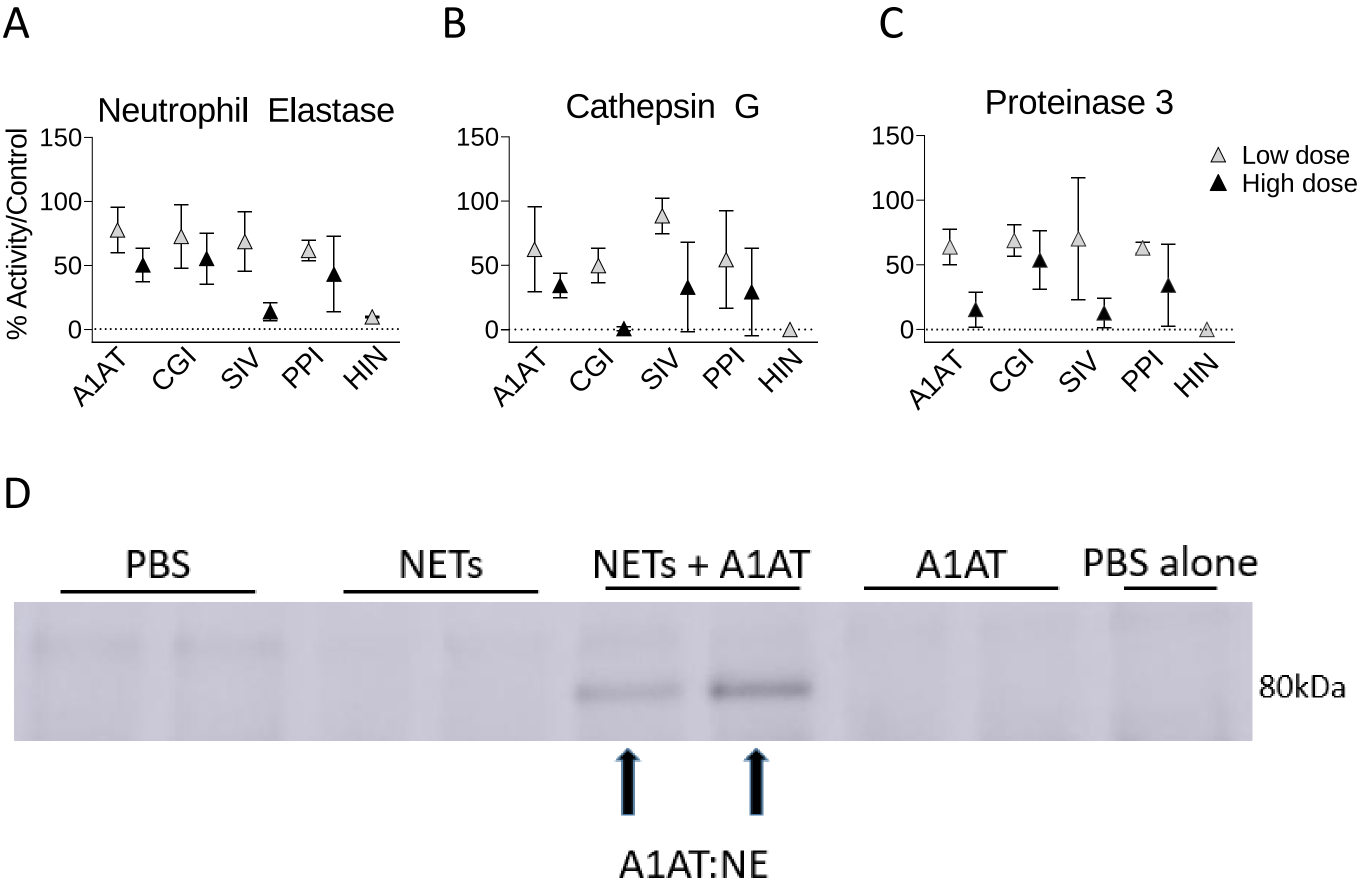


**Supplemental Figure 5. Protease inhibitors differentially reduce NET protease activities and can form complexes with NET proteases.** Isolated human NETs were incubated for 1h at 37°C with two concentrations of protease inhibitors: A1AT, cathepsin G inhibitor I (CGI), sivelestat an NE inhibitor (SIV), protease inhibitor cocktail III, a pan protease inhibitor (PPI), and heat inactivated NETs (HIN). Protease activity of **(A)** NE, **(B)** CG, and **(C)** PR3 were measured using fluorescent resonance energy transfer (FRET) substrates and are expressed as percent of the NETs alone control (experiments=3, NETs donors=3). **(D)** HBE were exposed to PBS, 5µg/ml NETs, NETs+100µg/ml A1AT or A1AT for 18h in triplicate wells. Immunoprecipitation (IP) was performed on HBE supernatants or PBS alone (no supernatant) using NE antibody to precipitate NE protein. Anti-A1AT primary antibody was used to visualize the 80k A1AT:NE complex band in the NETs+A1AT lanes (experiments=2, HBE donors=2, NET donors=2).

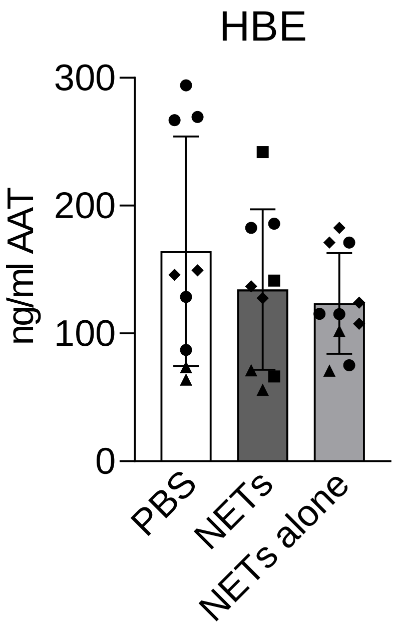


**Supplemental Figure 6. Human NETs contain A1AT and HBE secrete A1AT.** HBE grown at ALI were exposed to PBS control or 5µg/ml NETs in the apical compartment for 18h in duplicate to triplicate wells**.** HBE exposed to PBS or NETs secrete similar concentrations of A1AT into the apical supernatant. NETs alone were simultaneously incubated for 18 h without epithelial cells. Human A1AT concentrations were measured via ELISA. Each symbol represents NETs isolated from a different donor. Results were analyzed by one-way ANOVA. (experiments=3, HBE donors=3, NETs=3)

**Supplemental Table 1. NETs alter RNAs affecting apoptosis in human bronchial epithelia.** List of differentially expressed genes associated with apoptosis found by RNA sequencing of HBE exposed to 5μg/ml NETs compared to PBS for 18h. Data analyzed using IPA (p=1.07e-13, experiments=3, HBE donors=3, NET donors=3).

| Apoptosis | | | | | |
|---|---|---|---|---|---|
| Symbol | Fold Change | Symbol | Fold Change | Symbol | Fold Change |
| *IL1RL1* | 104.7046 | *HDAC9* | 5.019643 | *PBX1* | -2.89936 |
| *CD200* | 49.66355 | *VAV1* | 4.949888 | *PTGS1* | -3.01138 |
| *MMP9* | 49.35035 | *TIMP2* | 4.81805 | *LGR4* | -3.01154 |
| *SEMA7A* | 39.21162 | *PHLDA2* | 4.794256 | *PTTG1* | -3.09511 |
| *CEACAM1* | 22.26556 | *NOG* | 4.709203 | *WNT5A* | -3.24799 |
| *PLAUR* | 14.30711 | *PIK3AP1* | 4.528828 | *WNT3A* | -3.59439 |
| *RASSF2* | 13.18982 | *IL32* | 4.515641 | *FKBP5* | -3.85641 |
| *PLAU* | 10.91761 | *TLR2* | 4.487772 | *PIK3R1* | -3.97341 |
| *MMP10* | 9.987298 | *POLB* | 4.475649 | *ID1* | -3.97642 |
| *TIAM1* | 9.666262 | *SOX9* | 4.434872 | *JAG2* | -4.05528 |
| *GPR132* | 8.60728 | *CLCF1* | 4.37477 | *FGFR3* | -4.06072 |
| *GDF15* | 8.016345 | *EREG* | 4.200381 | *ID3* | -4.20011 |
| *TREM1* | 7.543849 | *CCND1* | 3.948307 | *VAV3* | -4.31285 |
| *IL1RN* | 7.038242 | *MAP4K4* | 3.876084 | *ANGPT1* | -4.32759 |
| *ELOVL4* | 6.957872 | *PTGS2* | 3.831697 | *HSD11B2* | -4.41233 |
| *DKK1* | 6.847217 | *CEACAM5* | 3.775852 | *SDC2* | -4.5023 |
| *NT5E* | 6.753523 | *TGFB1* | 3.726483 | *SNAI2* | -4.71535 |
| *GLIPR1* | 6.489568 | *KRT17* | 3.520094 | *CAV1* | -5.04368 |
| *PHLDA1* | 6.403348 | *RASD1* | 3.487722 | *TIMP3* | -5.35201 |
| *PLK3* | 6.377526 | *PIK3IP1* | 3.429813 | *NGFR* | -5.55712 |
| *OSBP2* | 5.996882 | *ITGA1* | 3.152443 | *MT3* | -6.51282 |
| *ADAM8* | 5.705564 | *F3* | 3.082455 | *APLN* | -7.14273 |
| *FST* | 5.428863 | *KIFC3* | 2.944804 | *GAS1* | -7.24106 |
| *EHD3* | 5.288924 | *TXNRD1* | 2.805253 | *ID2* | -7.33517 |
| *SPHK1* | 5.132752 | *SCD* | -2.69065 | *TP63* | -7.46925 |
| *MMP1* | 5.041066 | *PTHLH* | -2.78389 | *DLL1* | -9.40422 |
| *CCL5* | 5.024486 | *TRIM2* | -2.8052 | | |
| *IRAK2* | 5.024247 | *PDCD4* | -2.87253 | | |

**Supplemental Table 2. NETs alter epithelial cell junction RNAs.** List of functionally enriched RNAs for cellular components of anchored junctions and tight junctions found by RNA sequencing of HBE exposed to 5µg/ml NETs compared to PBS for 18h. Data analyzed using ToppGene (experiments=3, HBE donors=3, NET donors=3).

| Cell Junction Organization | |
|---|---|
| Symbol | Fold Change |
| *ACTB* | 2.89996469 |
| *KIFC3* | 2.944804134 |
| *CLDN4* | 3.207103512 |
| *TGFB1* | 3.726483192 |
| *MAP4K4* | 3.876084105 |
| *LAMB3* | 4.014288083 |
| *TLR2* | 4.487771729 |
| *ESAM* | 4.649265569 |
| *EPHB3* | 4.978859764 |
| *DKK1* | 6.847217229 |
| *LAMC2* | 8.371985395 |
| *TIAM1* | 9.666262173 |
| *ARHGAP22* | -5.618308079 |
| *CAV1* | -5.043682033 |
| *WNT4* | -4.864411143 |
| *CLDN8* | -4.838474606 |
| *SNAI2* | -4.715354949 |
| *PIK3R1* | -3.973412632 |
| *MTSS1* | -3.870409255 |
| *WNT3A* | -3.594393098 |
| *WNT5A* | -3.247994985 |
